# Discovery and Optimization of Small Molecule Inhibitors of the SLIT2/ROBO1 Protein-Protein Interaction Using DNA-Encoded Libraries

**DOI:** 10.64898/2026.02.21.707154

**Authors:** Nelson García-Vázquez, Shaoren Yuan, Moustafa T. Gabr

## Abstract

Graphical Abstract

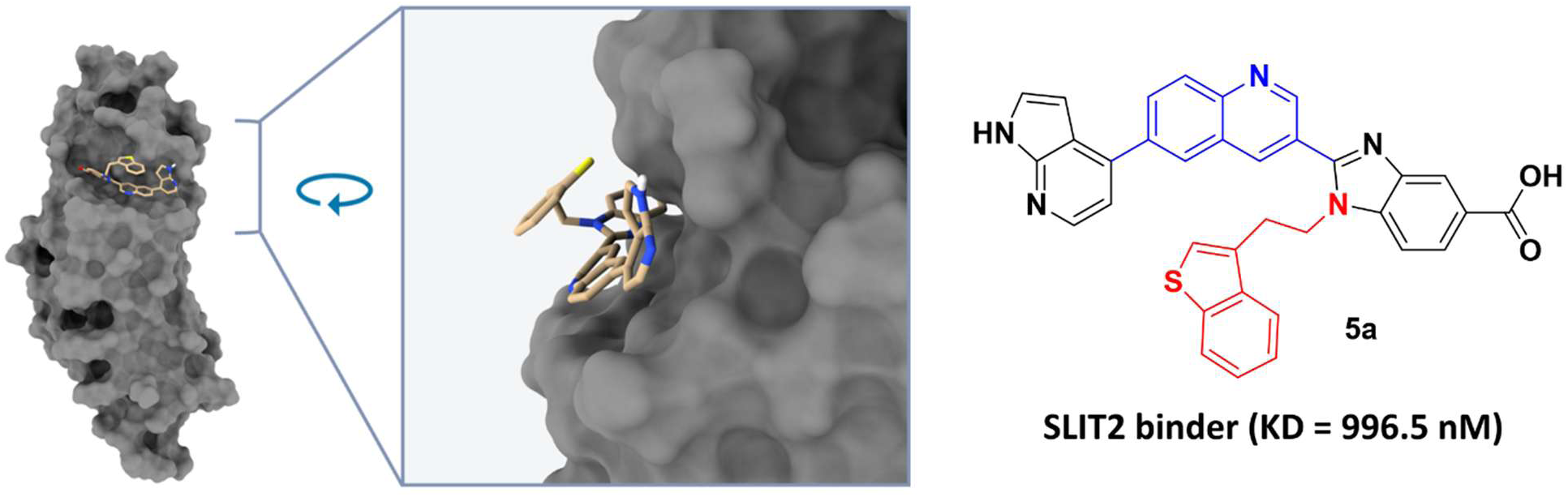

Protein-protein interactions (PPIs) mediated by extracellular ligands remain challenging targets for small molecule intervention due to their large and dynamic interfaces. The interaction between SLIT2 and its receptor ROBO1 plays a critical role in cell migration and tumor progression, yet remains largely unexplored. Here, we report the discovery and optimization of small molecule inhibitors of the SLIT2/ROBO1 interaction enabled by DNA-encoded library (DEL) screening. Affinity selection against SLIT2 identified four structurally diverse hit compounds, which were subsequently validated using orthogonal biophysical assays. Among these, one hit exhibited measurable SLIT2 binding and functional inhibition of the SLIT2/ROBO1 interaction in a time-resolved FRET assay. Guided by physicochemical considerations, a solubility-optimized analog was designed, resulting in a ∼50-fold improvement in binding affinity and an ∼9-fold enhancement in functional potency. Molecular dynamics simulations and induced-fit docking revealed a stable binding mode within the SLIT2 LRR2 domain and suggested that a benzothiophene substituent was dispensable for target engagement. Fragment-based experimental validation confirmed this prediction, leading to the identification of a minimal azaindole-based pharmacophore that retained nanomolar binding affinity. Collectively, this study demonstrates how DEL-enabled hit discovery combined with rational optimization and fragment deconstruction can yield potent small molecule modulators of a challenging extracellular PPI, providing a foundation for further development of SLIT2/ROBO1 pathway inhibitors.

## 1. Introduction

Slit guidance ligands (SLITs) constitute a family of secreted extracellular proteins that modulate spatial cell organization during development through interactions with roundabout (ROBO) receptors.^1^ In mammals, three SLIT isoforms (SLIT1, SLIT2, and SLIT3) engage ROBO1 and ROBO2 via binding of the second leucine-rich repeat (D2) domain of SLITs to the Ig1 domain of ROBO receptors.^2,3^ Among these, SLIT2-mediated signaling has emerged as a central regulator of diverse biological processes extending well beyond developmental guidance. Accumulating evidence implicates the SLIT2/ROBO axis in organogenesis, tissue homeostasis, and cancer progression, where it modulates not only directional cell migration but also proliferation, apoptosis, adhesion, angiogenesis, and fibrotic remodeling. In particular, SLIT2-driven recruitment of adaptor proteins to the ROBO cytoplasmic tail enables dynamic cytoskeletal remodeling, thereby influencing cell motility, growth, and survival across multiple physiological and pathological contexts.^3–5^ Additionally, SLIT2/ROBO signaling has been linked to liver fibrosis through activation of PI3K/Akt pathways, underscoring its broad systemic relevance.

Within the nervous system, SLIT2/ROBO signaling plays a foundational role in axonal repulsion,^6,7^ neuronal migration,^8^ and axon pathfinding.^9^ Beyond neurobiology, SLIT ligands exert complex and cell type-specific effects in the immune system, acting as chemoattractants for neutrophils while simultaneously repelling lymphocytes and dendritic cells.^10–12^ In macrophages, SLIT signaling suppresses macropinocytosis and cytotoxic polarization,^13^ whereas in endothelial cells, SLIT2/ROBO activation promotes angiogenic programs by directing tip cell migration and polarization during vascular development in retinal and skeletal tissues.^14–16^ These pleiotropic functions extend into oncology, where SLIT2 has been shown to enhance tumor angiogenesis,^17^ facilitate tumor cell migration,^18–20^ promote metastatic dissemination,^21^ and contribute to resistance against anticancer therapies.^22^ Such tumor-promoting activities have been reported in colorectal cancer, pancreatic cancer, and osteosarcoma, among others. Conversely, SLIT2/ROBO signaling has also been associated with tumor-suppressive effects in lung and breast cancers,^23–25^ highlighting the context-dependent nature of this pathway. In glioblastoma (GBM), particularly, both inhibitory^26–28^ and pro-tumorigenic^29,30^ roles have been described, emphasizing the complexity of SLIT2/ROBO signaling within distinct cellular and microenvironmental settings.

More recently, SLIT2/ROBO signaling has been identified as a critical regulator of immune evasion within the GBM tumor microenvironment (TME).^31^ Elevated SLIT2 expression in patient-derived samples and genetically engineered mouse models was shown to drive the accumulation of immunosuppressive tumor-associated macrophages (TAMs) and induce aberrant vascular architecture.^31^ Genetic silencing of SLIT2 in glioma cells, as well as systemic blockade using the SLIT2-trapping biologic ROBO1Fc, effectively prevented TAM polarization toward tumor-supportive phenotypes and suppressed angiogenic gene expression programs.^31^ These interventions resulted in improved tumor vessel normalization and significantly enhanced responses to both chemotherapy and immunotherapy in GBM models.^31^ Notably, inhibition of SLIT2 produced more pronounced effects on angiogenesis and T-cell-mediated immune responses than several previously investigated strategies aimed at TAM modulation within the GBM TME.^31^

Despite the growing therapeutic interest in the SLIT2/ROBO pathway, clinical translation has thus far been limited. The only completed clinical trial involving a SLIT2/ROBO-targeting biotherapeutic (PF-06730512) was conducted in patients with focal segmental glomerulosclerosis (FSGS) but was discontinued in 2023 due to insufficient efficacy at tolerable dose levels.^32,33^ In parallel, three ongoing clinical studies (NCT03940820, NCT03941457, and NCT03931720) are evaluating chimeric antigen receptor-natural killer (CAR-NK) cell therapies targeting ROBO1 in solid tumors, reflecting continued interest in this axis as a therapeutic target.

While biologic modalities such as receptor traps and monoclonal antibodies have demonstrated proof-of-concept activity, small molecule inhibitors offer several potential advantages over protein-based therapies, including reduced immunogenicity, improved tissue penetration, and greater flexibility in managing adverse events due to favorable pharmacokinetic profiles.^34–37^ Oral bioavailability and shorter systemic exposure further reduce the risk of prolonged on-target immune-related toxicities, positioning small molecules as an attractive alternative for targeting immune-modulatory pathways.^38,39^ Despite these advantages and the compelling biological rationale, no small molecule inhibitors of SLIT2/ROBO1 signaling have been reported to date, representing a significant unmet need in drug discovery. Challenges inherent to biologic therapies, including manufacturing complexity, limited tumor penetration, and potential immune activation, further underscore the urgency of developing small molecule modulators capable of disrupting SLIT2-driven tumor progression and immune suppression.^34–39^ The identification of such compounds could enable scalable, cost-effective therapeutic strategies for GBM and other malignancies in which SLIT2/ROBO signaling plays a pathogenic role.

DNA-encoded library (DEL) technology has emerged as a highly efficient strategy for the identification of small molecules that reversibly engage protein targets, providing access to chemical space that far exceeds the scope of conventional high-throughput screening (HTS) approaches.^40–42^ DEL platforms rely on libraries of DNA-tagged small molecules, in which each compound is covalently linked to a unique oligonucleotide barcode that records its synthetic history. ^40–42^ Library construction typically involves iterative cycles of split-and-pool synthesis, where sequential chemical transformations are coupled with stepwise DNA barcode elongation.^40,41^ Through this modular design, the combinatorial use of even a modest number of building blocks can generate libraries containing hundreds of millions to billions of distinct compounds; for example, three synthetic cycles incorporating approximately 1,000 building blocks per step can theoretically yield 10^9^ unique entities. The resulting libraries enable pooled affinity-based screening experiments that require only minimal quantities of purified target protein, substantially improving throughput and cost efficiency relative to traditional screening paradigms.^40,41^ During selection, DEL compounds are incubated with immobilized target proteins, allowing high-affinity binders to be retained while non-interacting species are removed through stringent washing. Enriched ligands are subsequently recovered and identified by high-throughput DNA sequencing, in which the attached barcodes function as molecular identifiers, enabling rapid and unbiased hit discovery from extremely large chemical collections.^40,41^

In the present study, we applied DEL screening to identify small molecule binders of SLIT2, followed by biophysical and functional validation, medicinal chemistry optimization, and fragment-based deconstruction to define a minimal pharmacophore capable of disrupting the SLIT2/ROBO1 protein-protein interaction. This work establishes a tractable small molecule strategy for targeting the SLIT2/ROBO1 signaling axis, addressing a critical gap in therapeutic approaches for modulating this pathway.

## 2. Results and discussion

### 2.1. Identification of SLIT2-binding compounds from DNA-encoded library screening

DEL screening against SLIT2 was performed using a large, highly diverse library comprising approximately 1.4 billion compounds (AIphaMa DEL screening kit), enabling extensive coverage of chemical space under affinity-based selection conditions. Following three iterative rounds of target panning and enrichment, analysis of next-generation sequencing data revealed a limited number of compounds displaying clear enrichment relative to no-target control selections. These enriched species exhibited well-defined structure-enrichment relationships and reproducible copy number increases across selection rounds, consistent with specific target engagement rather than nonspecific binding. From this analysis, four structurally distinct small molecule hits (**NS-01**-**NS-04**) were prioritized for off-DNA resynthesis and experimental validation (**Figure 1**). The identification of multiple chemotypes from a single DEL campaign highlights both the robustness of the selection strategy and the presence of druggable binding features on SLIT2 amenable to small-molecule recognition.

**Figure 1.**
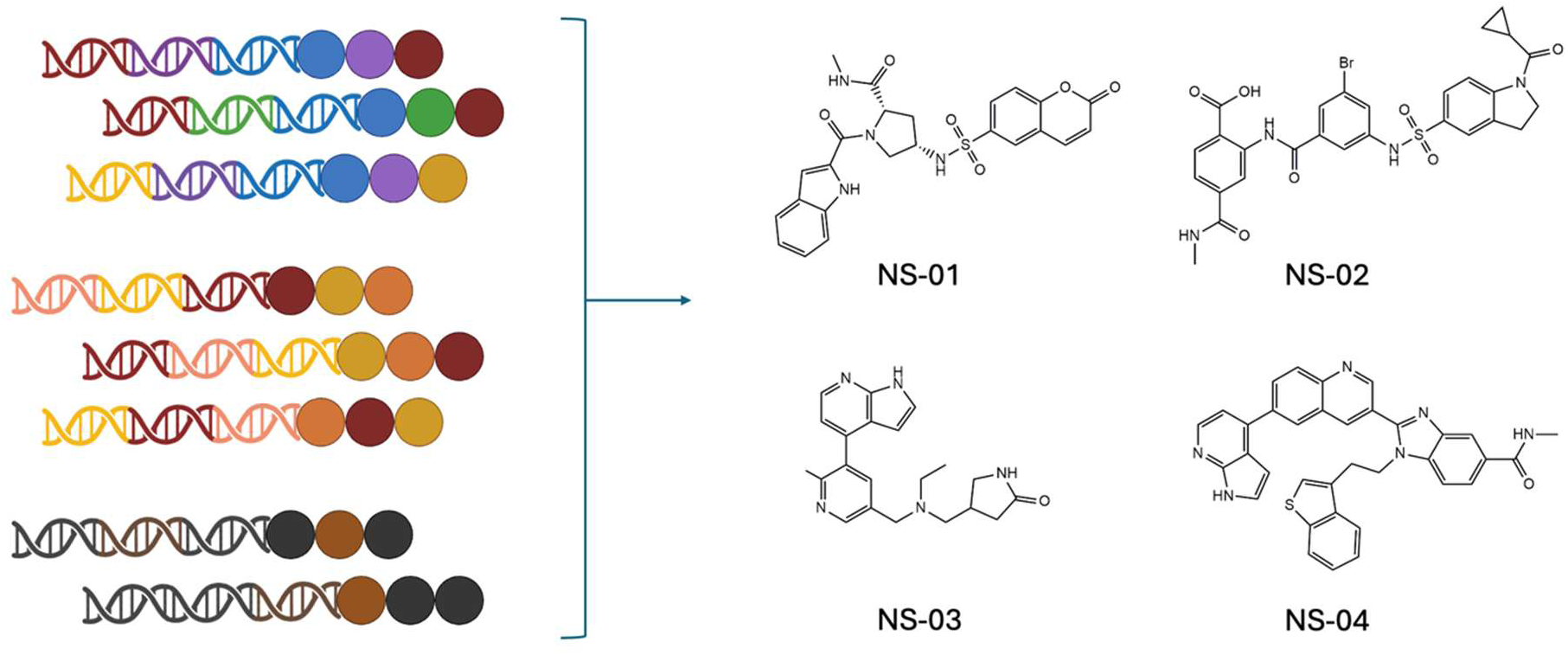
DEL screening identifies four structurally diverse SLIT2-binding hit compounds. Schematic representation of DNA-encoded library (DEL) screening strategy (left) and chemical structures of the four hit compounds identified from the AlphaMa DEL screen (right). Compounds **NS-01**, **NS-02**, **NS-03**, and **NS-04** represent distinct chemical scaffolds selected for further characterization based on enrichment in the SLIT2 affinity selection.

To orthogonally validate DEL-derived hits and quantify their binding affinities, we employed two independent biophysical assays, Temperature-Related Intensity Change (TRIC) and Spectral Shift. Among the four resynthesized compounds, **NS-04** displayed the most pronounced and reproducible binding to SLIT2, exhibiting a dissociation constant (Kd) of 46.5 μM by TRIC (**Figure 2D**). **NS-02** emerged as the second most potent binder with a Kd of 71.5 μM, while **NS-01** and **NS-03** failed to produce convincing dose-response curves indicative of specific binding (**Figure 2A, 2C**). These results established **NS-04** as the most promising DEL-derived binder for subsequent functional evaluation and medicinal chemistry optimization.

**Figure 2.**
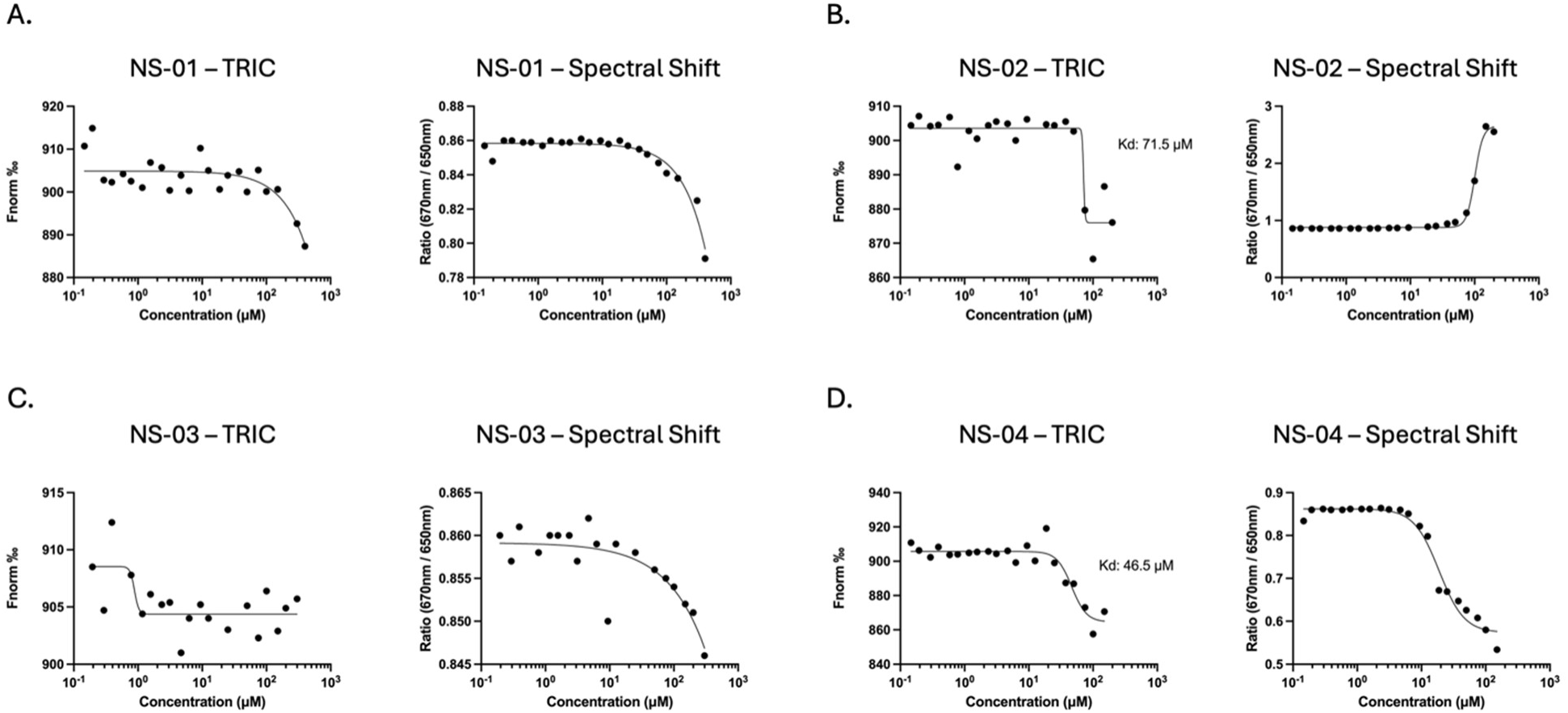
Biophysical characterization of DEL hits by TRIC and Spectral Shift. Dose-response curves for SLIT2 binding by TRIC (left panels) and Spectral Shift (right panels) for (**A**) **NS-01**, (**B**) **NS-02**, (**C**) **NS-03**, and (**D**) **NS-04**. **NS-04** exhibited the most favorable binding profile with a Kd of 46.5 μM by TRIC, while NS-02 showed a Kd of 71.5 μM. NS-01 and NS-03 did not produce convincing binding curves. Data points represent individual measurements.

Following confirmation of direct SLIT2 binding, we next evaluated whether the DEL-derived compounds could functionally disrupt the SLIT2/ROBO1 PPI. Functional activity was assessed using a time-resolved fluorescence resonance energy transfer (TR-FRET) assay developed in-house, in which ROBO1-Fc and His-tagged SLIT2 are detected via terbium- and D2-conjugated antibodies, respectively (**Figure 3A**). Under these conditions, **NS-04** was the only compound to exhibit concentration-dependent inhibition of the SLIT2/ROBO1 interaction, with an EC_50_ value of 79.3 μM and a maximal inhibition of approximately 58% at the highest concentration tested (**Figure 3E**). In contrast, **NS-01**, **NS-02**, and **NS-03** displayed no appreciable inhibitory activity in this assay (**Figure 3B-D**), indicating that SLIT2 binding alone was insufficient to confer functional PPI disruption. These data further prioritized **NS-04** as the only DEL-derived hit exhibiting both biophysical binding and functional antagonism of the SLIT2/ROBO1 interaction.

**Figure 3.**
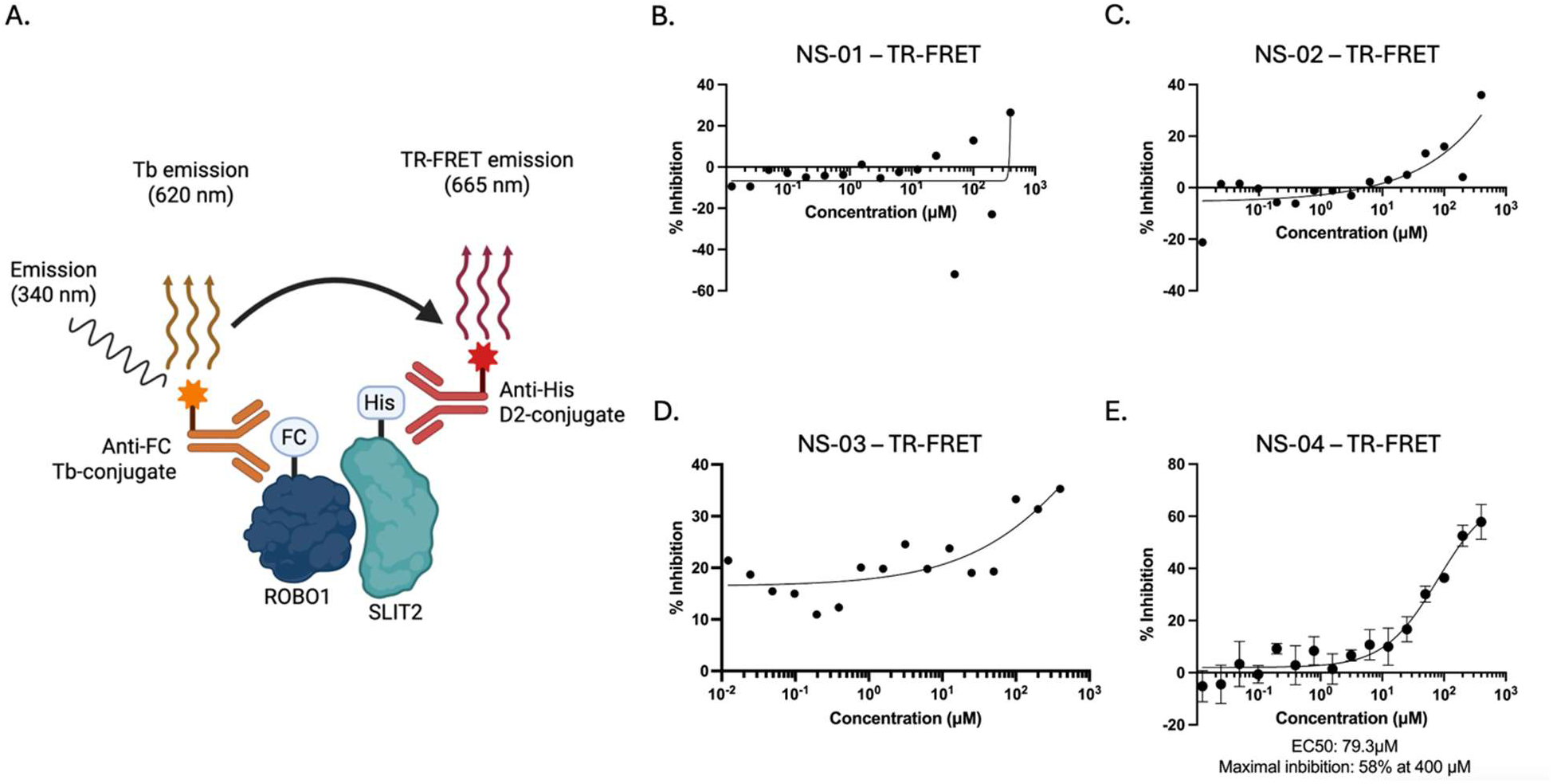
TR-FRET assay for SLIT2/ROBO1 protein-protein interaction inhibition. (**A**) Schematic of the TR-FRET assay design. ROBO1-Fc is detected via anti-Fc terbium (Tb) conjugate (emission 620 nm), and His-tagged SLIT2 is detected via anti-His D2 conjugate. Upon excitation at 340 nm, energy transfer between bound ROBO1 and SLIT2 produces TR-FRET emission at 665 nm. (**B-E**) Dose-response curves showing percent inhibition of the SLIT2/ROBO1 interaction for (**B**) **NS-01**, (**C**) **NS-02**, (**D**) **NS-03**, and (**E**) **NS-04**. Only **NS-04** demonstrated appreciable inhibition with an EC_50_ of 79.3 μM and maximal inhibition of 58% at 400 μM. Error bars represent standard deviation.

### 2.2. Development of an improved SLIT2 binder through solubility optimization

During preliminary handling and assay preparation, **NS-04** exhibited limited aqueous solubility at higher concentrations, accompanied by evidence of aggregation, indicating an unfavorable physicochemical profile. This behavior was not observed for **NS-02**, suggesting that solubility, rather than intrinsic binding capability, represented a key limitation for **NS-04**. Despite this liability, the combined biophysical and functional activity of **NS-04** supported its advancement for structure-guided optimization. To address the solubility limitations observed for **NS-04** while enabling preliminary structure-activity relationship (SAR) exploration, a convergent synthetic strategy was developed that allowed late-stage diversification of the N-substituent and aromatic heterocycle (**Scheme 1**). The route commenced with nucleophilic aromatic substitution of ethyl 4-fluoro-3-nitrobenzoate (**1**) with substituted amines, followed by nitro reduction to afford an aniline intermediate (**3**) amenable to heterocycle formation. Condensation with aryl aldehydes enabled construction of the benzimidazole core, providing access to ester intermediates (**4**) that could be readily hydrolyzed to the corresponding carboxylic acids (**5**). This modular sequence facilitated efficient conversion of the parent amide scaffold into the corresponding acid analogue (**5**a, **5b**, and **5c**), as well as preparation of related amide derivatives (**6b** and **6c**) with minimal synthetic re-optimization (**Scheme 1**).

Importantly, the introduction of the carboxylic acid functionality was designed to improve aqueous solubility while preserving the core heterocyclic framework implicated in SLIT2 binding. The same synthetic pathway also enabled rapid access to amide analogues through late-stage coupling, allowing direct comparison of acid and amide variants within a consistent structural context. This flexibility proved essential for subsequent fragment-based deconstruction and SAR analysis, as it enabled systematic evaluation of substituent contributions to binding affinity and functional activity while maintaining control over physicochemical properties.

**Scheme 1.**
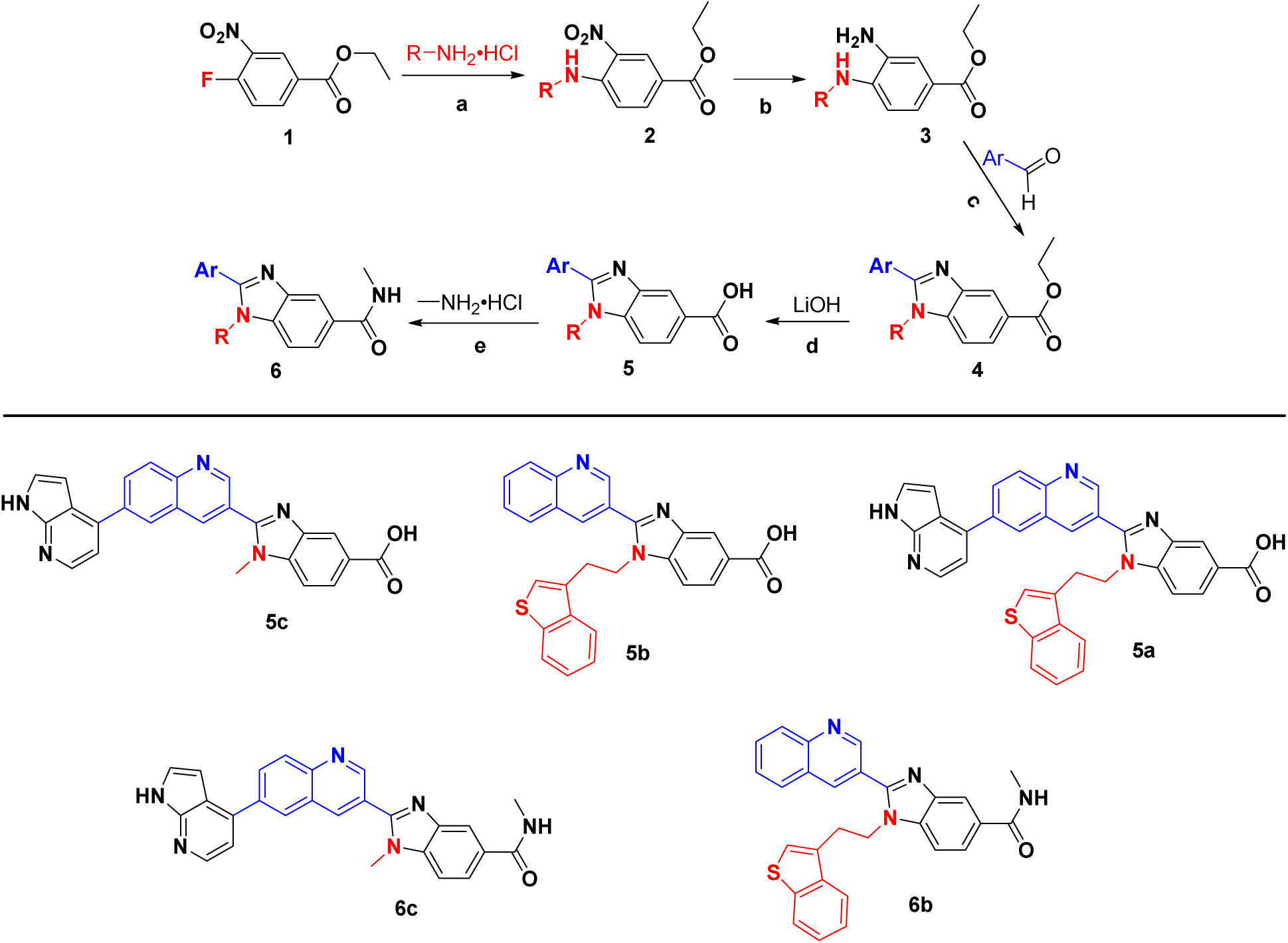
Synthetic route enabling solubility optimization and SAR exploration of SLIT2 binders (top) and chemical structures of the synthesized compounds (bottom). Reagents and conditions: (a) R–NH₂·HCl, base; (b) reduction of the nitro group to the corresponding aniline; (c) condensation with aryl aldehydes to form the benzimidazole core; (d) ester hydrolysis to afford the corresponding carboxylic acid (compound 5); (e) conversion to the amide analogue (compound 6). This modular synthetic strategy enabled late-stage diversification and direct comparison of acid and amide derivatives while preserving the core heterocyclic scaffold.

Biophysical evaluation of **5a** (**Figure 4A**) revealed a marked enhancement in SLIT2 binding affinity, with TRIC analysis yielding a Kd of 996.5 ± 138.0 nM and Spectral Shift measurements providing a consistent Kd of 1.25 ± 0.26 μM (**Figure 4B,C**). Relative to the parent compound **NS-04**, this corresponds to an approximately 47-fold improvement in binding affinity as determined by TRIC. Importantly, these gains translated into substantially improved functional activity, as **5a** inhibited the SLIT2/ROBO1 interaction in the TR-FRET assay with an EC_50_ of 9.01 ± 0.95 μM and a maximal inhibition of 77% at 50 μM (**Figure 4D**). Compared with **NS-04**, this represents an 8.8-fold increase in functional potency together with a notable increase in maximal inhibitory efficacy.

**Figure 4.**
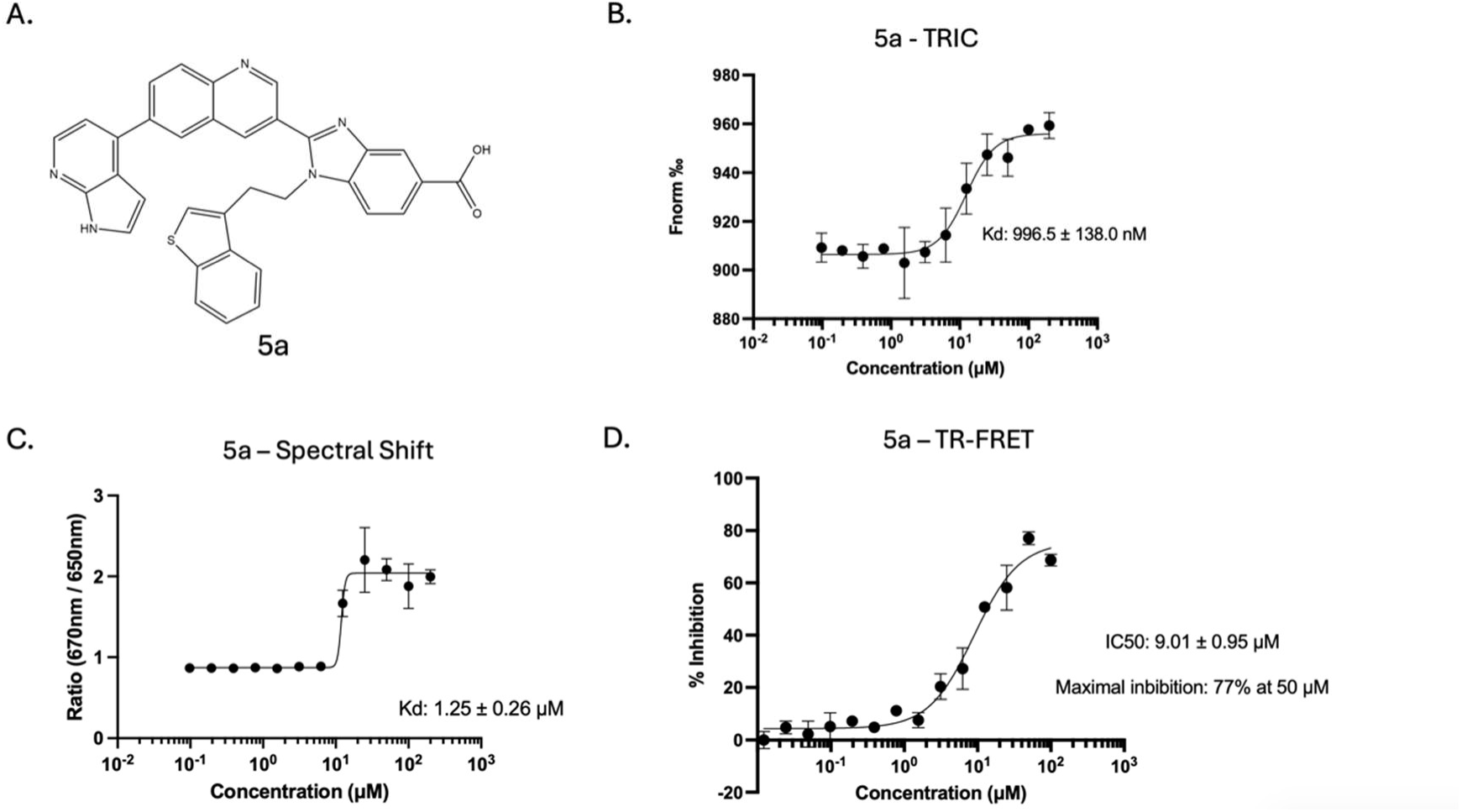
Compound 5a exhibits improved binding affinity and functional activity. (**A**) Chemical structure of compound **5a**, designed by replacing the amide of **NS-04** with a carboxylic acid to improve solubility. (**B**) TRIC dose-response curve yielding a Kd of 996.5 ± 138.0 nM. (**C**) Spectral Shift dose-response curve yielding a Kd of 1.25 ± 0.26 μM. (**D**) TR-FRET inhibition assay demonstrating an EC_50_ of 9.01 ± 0.95 μM and maximal inhibition of 77% at 50 μM. Error bars represent standard deviation.

### 2.3. Molecular dynamics simulations reveal the binding mode and identify non-essential structural features

To gain mechanistic insight into the binding mode of **5a** and to inform subsequent structure-guided optimization, molecular dynamics (MD) simulations were performed using the SLIT2 leucine-rich repeat 2 (LRR2) domain. To minimize bias associated with predefined docking poses and to enhance sampling of productive binding events, the system was solvated with six copies of **5a**. Triplicate 100 ns simulations were conducted, and ligand-protein contacts were quantified across the trajectories. Residues exhibiting sustained interactions were mapped and visualized using a viridis color scale, where purple indicates negligible contact and yellow denotes persistent engagement throughout the simulation window (**Figure 5A**).

**Figure 5.**
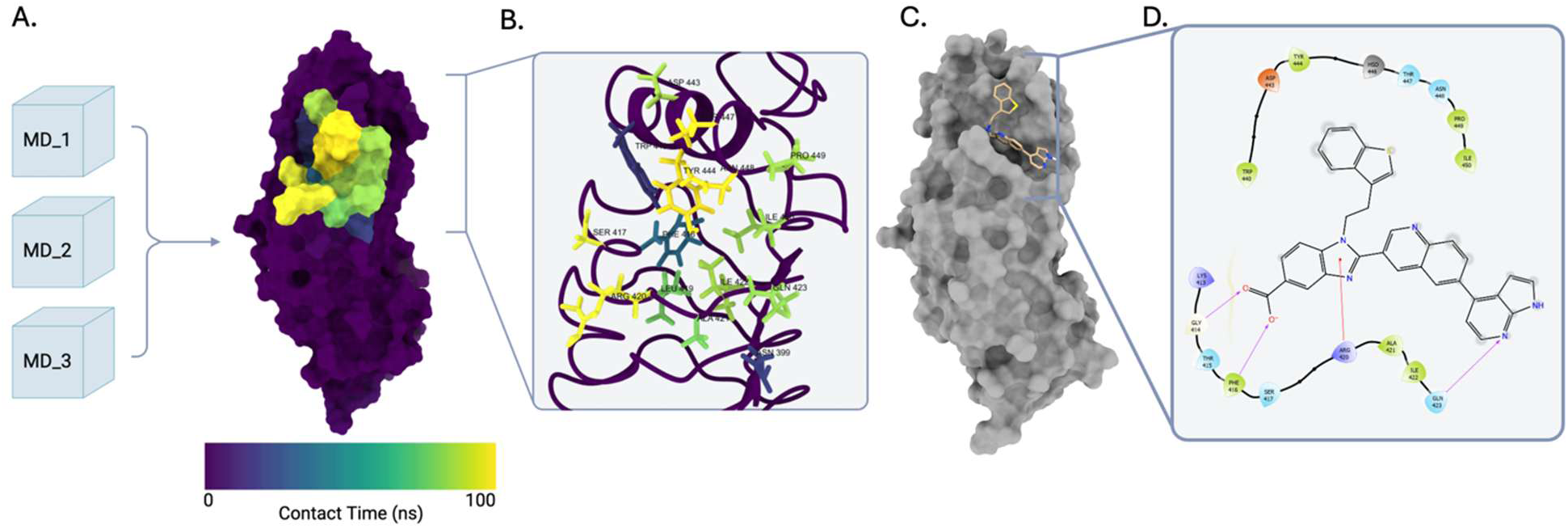
Molecular dynamics simulations and induced fit docking identify the SLIT2 binding site. (**A**) Surface representation of the SLIT2 LRR2 domain colored by ligand contact time from triplicate 100 ns MD simulations. Color scale ranges from purple (0 ns contact) to yellow (100 ns contact). (**B**) Ribbon diagram highlighting residues involved in ligand contacts: TYR444, THR447, ASN448, ARG420, SER417, ASP443, ILE422, PRO449, ILE450, GLN423, ALA421, LEU419, PHE416, ASN399, and TRP440. (**C**) Surface representation showing the top pose from induced fit docking (IFD) of **5a** within the predicted binding site. (**D**) Two-dimensional interaction diagram illustrating key contacts between **5a** and SLIT2 binding site residues.

Analysis of the combined trajectories revealed a reproducible cluster of SLIT2 residues involved in ligand recognition, including TYR444, THR447, ASN448, ARG420, SER417, ASP443, ILE422, PRO449, ILE450, GLN423, ALA421, LEU419, PHE416, ASN399, and TRP440 (**Figure 5B**). The convergence of these contact patterns across independent simulations supports the presence of a defined ligand-interaction region within the LRR2 domain and provided a structural framework for subsequent docking refinement and fragment-based deconstruction.

To refine the predicted binding pose, we performed induced fit docking (IFD) of **5a** within the binding site defined by the MD contact analysis. This approach allows for conformational flexibility of both the ligand and the protein side chains, yielding poses that account for induced fit effects upon ligand binding (**Figure 5C, 5D**). The top-ranked IFD pose was subsequently subjected to additional MD simulations to assess pose stability. Throughout these simulations, **5a** exhibited only minor rearrangements within the binding pocket while maintaining a stable overall orientation (**Figure 6A, 6B**). Notably, the benzothiophene moiety remained predominantly solvent-exposed in the stable binding pose, suggesting that this structural element may not be essential for SLIT2 recognition (**Figure 6C**).

**Figure 6.**
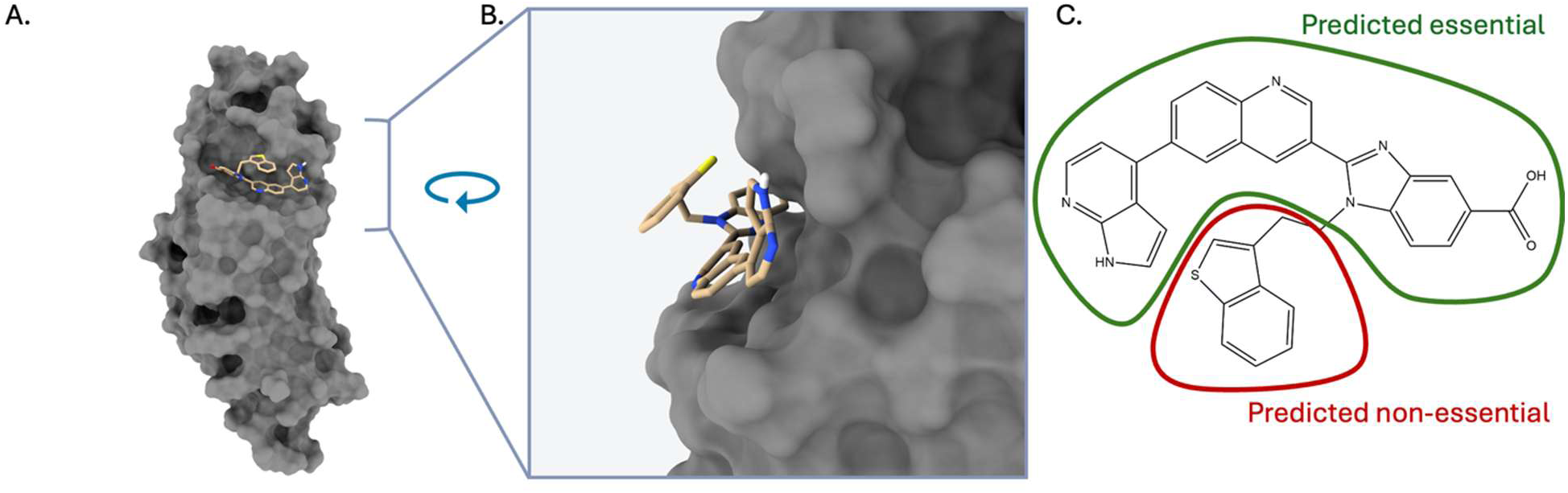
MD simulation of the top IFD pose reveals a stable binding mode with solvent-exposed benzothiophene. (**A**) Surface representation of SLIT2 with **5a** in the representative stable pose from MD simulation initiated from the top IFD pose. (**B**) Magnified view of the binding pocket showing **5a** orientation, with the benzothiophene moiety projecting toward solvent. (**C**) Chemical structure of **5a** annotated to indicate regions predicted to be essential for binding (green outline) and non-essential (red outline: benzothiophene moiety).

### 2.4. Fragment-based validation confirms dispensability of the benzothiophene moiety

To directly test the hypothesis derived from molecular dynamics simulations that the benzothiophene moiety is dispensable for SLIT2 binding, we designed and synthesized two complementary fragment derivatives of **5a**. Compound **5b** retains the benzothiophene substituent while lacking the azaindole core, whereas compound **5c** preserves the azaindole moiety with removal of the benzothiophene group (**Figure 7A,C**). Biophysical evaluation demonstrated that 5b failed to exhibit measurable binding to SLIT2 in either TRIC or Spectral Shift assays (**Figure 7B**), indicating that the benzothiophene alone is insufficient for target engagement. In contrast, 5c retained high-affinity binding, with dissociation constants of 898 nM by TRIC and 822 nM by Spectral Shift (**Figure 7D**). Notably, 5c displayed a modest improvement in affinity relative to the parent compound 5a (1.1-fold by TRIC and 1.5-fold by Spectral Shift), providing experimental confirmation that the benzothiophene moiety contributes minimally to binding and validating the azaindole-based core as the primary pharmacophoric element.

**Figure 7.**
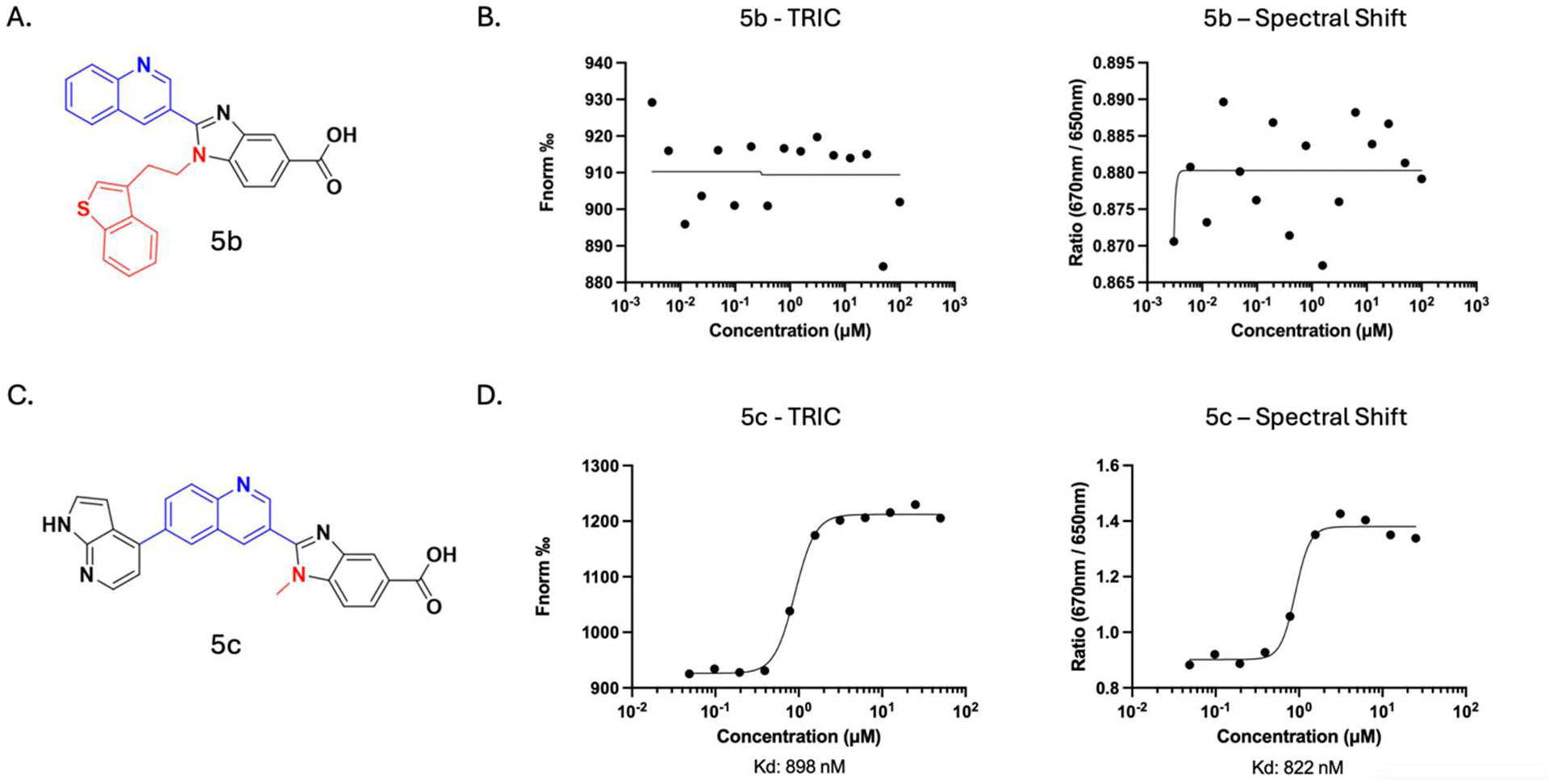
Fragment analysis of carboxylic acid derivatives confirms dispensability of the benzothiophene. (**A**) Chemical structure of fragment **5b**, retaining the benzothiophene while lacking the azaindole. (**B**) TRIC and Spectral Shift dose-response curves for **5b** showing no measurable binding to SLIT2. (**C**) Chemical structure of fragment **5c**, retaining the azaindole while lacking the benzothiophene. (**D**) TRIC (Kd = 898 nM) and Spectral Shift (Kd = 822 nM) dose-response curves demonstrating retained binding affinity for **5c**.

This trend was corroborated using the corresponding amide-containing fragments, **6b** and **6c** (**Figure 8A, 8C**). Compound **6b**, bearing the benzothiophene, showed no evidence of SLIT2 binding (**Figure 8B**), whereas **6c**, containing the azaindole core, demonstrated measurable affinity with a Kd of 22.3 μM by TRIC and 26.6 μM by Spectral Shift (**Figure 8D**). This represents a 2.1-fold improvement over the original DEL hit NS-04 (TRIC Kd 46.5 μM versus 22.3 μM), achieved despite the removal of a substantial portion of the molecular scaffold. These findings validate our computational predictions and establish that the azaindole-benzimidazole core constitutes the minimal pharmacophore required for SLIT2 recognition, while the benzothiophene moiety is dispensable and may be replaced with alternative substituents in future optimization campaigns.

**Figure 8.**
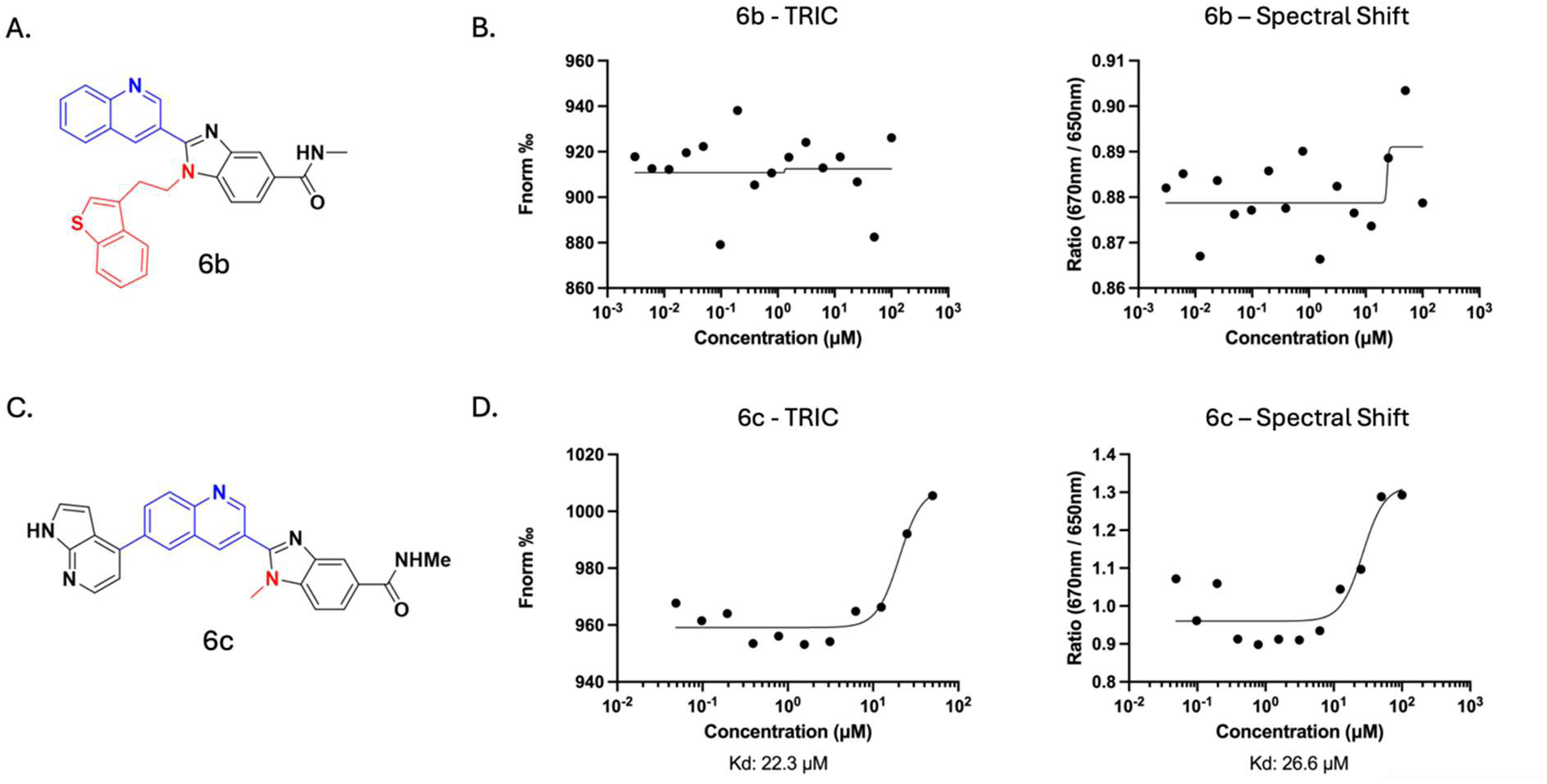
Fragment analysis of amide derivatives corroborates the minimal pharmacophore. (**A**) Chemical structure of fragment **6b**, the amide analog of **5b**, retaining the benzothiophene. Color coding as in Figure 7. (**B**) TRIC and Spectral Shift dose-response curves for **6b** showing no measurable binding to SLIT2. (**C**) Chemical structure of fragment **6c**, the amide analog of **5c**, retaining the azaindole core. (**D**) TRIC (Kd = 22.3 μM) and Spectral Shift (Kd = 26.6 μM) dose-response curves demonstrating **6c** retains binding with a 2.1-fold improvement over the parent compound NS-04.

## 3. Conclusion

In this study, we report a DEL-enabled strategy for the discovery and optimization of small-molecule inhibitors targeting the SLIT2/ROBO1 protein–protein interaction. Starting from DEL-derived hit compounds, we established direct SLIT2 binding and functional disruption of the SLIT2/ROBO1 interaction using complementary biophysical and TR-FRET assays. Rational optimization guided by physicochemical considerations led to a substantial improvement in both binding affinity and functional potency, while molecular dynamics simulations provided mechanistic insight into ligand engagement within the SLIT2 LRR2 domain. Importantly, fragment-based experimental validation confirmed computational predictions and enabled identification of a minimal azaindole-based pharmacophore sufficient for high-affinity SLIT2 recognition.

Collectively, these findings demonstrate that extracellular PPIs such as SLIT2/ROBO1, which have historically been addressed using biologic modalities, can be rendered tractable to small-molecule intervention through DEL-enabled discovery and structure-guided medicinal chemistry. The chemical framework and mechanistic insights established here provide a foundation for further optimization toward more potent and selective SLIT2/ROBO1 inhibitors and offer a generalizable approach for targeting challenging extracellular signaling interactions.

## 4. EXPERIMENTAL

### 4.1. Materials

hSLIT2-His was obtained from Sino Biological (Cat. No. 11967-H08H). hROBO1-Fc was obtained from Sino Biological (Cat. No. 30073-H02H). RED-tris-NTA 2nd generation dye was obtained from NanoTemper (Cat. No. MO-L018). Terbium-conjugated anti-human IgG polyclonal antibody (Cat. No. 61HFCTAF), d2-conjugated anti-His monoclonal antibody (Cat. No. 61HISDLF), and PPI Tb detection buffer (Cat. No. 61DB10RDF) were obtained from Cisbio. White 384-well plates were obtained from Greiner (Cat. No. 784075). HEPES solution (1 M in H₂O) was obtained from Millipore Sigma (Cat. No. 3264). Tween 20 (Ultrapure) was obtained from Thermo Scientific Chemicals (Cat. No. J20605.AP). Dimethyl sulfoxide (ACS, 99.9% min) was obtained from Thermo Scientific Chemicals (Cat. No. 036480.K2).

### 4.2. Protein immobilization tests

Prior to DEL screenings, protein immobilization tests were performed to confirm SLIT2 can bind to the magnetic beads through his tag. 10 ul of Hispur^TM^ Ni-NTA magnetic beads (Thermo Fisher Scientific, 88832) were incubated with 5 µg each target protein. The composition of the incubation buffer was 1x PBS, 10 mM imidazole, 0.1 mg/ml sssDNA, 0.005% Tween-20, pH 7.4. After 30 min incubation at RT, the supernatant was collected, and the beads were resuspended and washed by 100 µl of the incubation buffer. The supernatant was collected as wash, and the beads were resuspended by 100 µl of the incubation buffer. The bead-suspensions were then heated at 95 °C for 10 min, and the supernatant was collected as heated elution. The heated beads were again resuspended in 100 µl of the incubation buffer. 10 µl of each sample was loaded to a 5%/12% Tris-HCl SDS-PAGE. The gel was run for 150 V for 1 h followed by staining with Coomassie Blue for 1 h and destaining in distilled water overnight.

### 4.3. DEL screening against the target proteins

A 1.4 billion DNA-Encoded Chemical Library Kit (AlphaMa SDEL Pubkit) was purchased from AlphaMa Biotechnology, Suzhou. One kit afforded one screening condition against either the target or blank control bead as a no-target screening (NTC). DEL screenings against SLIT2 was performed by panning 10 µg target on 20 µl Hispur^TM^ Ni-NTA with the DNA-Encoded Chemical Libraries. Three rounds of affinity screening were performed in each condition, including protein immobilization, DEL panning, wash steps, and heated elution, following manufacturer’s instructions. Heated elution samples from the third round of affinity screening were subjected to the Next-Generation Sequencing to reveal the identity of the bound compounds under each screening condition. Utilizing AlphaMa’s decoding program (not disclosed), the fastq files were converted into chemical identities and the corresponding copies in each screening condition. The data were visualized by DataWarrior program. Each dot in the 2D and 3D plots represents a full molecule with each axis representing the BB index. The molecules with clear structure-enrichment relationship and high copies were considered as promising binders of the target and subjected to off-DNA hit resynthesis and validation.

### 4.4. Binding Affinity Measurements

Binding affinity measurements were performed on a Monolith X instrument (NanoTemper, Munich, Germany). Compounds were prepared as 12-point, 2-fold dilution series starting from 200 μM, followed by incubation with 50 nM hSLIT2-His labeled with RED-tris-NTA 2nd generation dye for 15 min at room temperature. Temperature-Related Intensity Change (TRIC) and Spectral Shift measurements were performed at 100% laser excitation and medium MST power. Kd values were calculated from Fnorm values obtained from three independent measurements using GraphPad Prism 10.

### 4.5. TR-FRET Protein-Protein Interaction Inhibition Assay

Inhibition of the SLIT2/ROBO1 interaction was evaluated using a TR-FRET assay. Recombinant human SLIT2 bearing a C-terminal His-tag (Sino Biological Cat. No. 11967-H08H) and the extracellular domain of ROBO1 fused to human IgG1 Fc (Sino Biological Cat. No. 30073-H02H) were employed. Detection was achieved using a terbium-conjugated anti-human IgG polyclonal antibody (Cisbio Cat. No. 61HFCTAF) and a d2-conjugated anti-His monoclonal antibody (Cisbio Cat. No. 61HISDLF).

Assay mixtures were prepared with final concentrations of 5 nM for both SLIT2 and ROBO1, 0.25 nM for the Tb-conjugated antibody, and 2.5 nM for the d2-conjugated antibody. All reagents were diluted in PPI Tb detection buffer (Cisbio Cat. No. 61DB10RDF). Compounds (final concentration: 400 μM in 3% DMSO) were dispensed into white 384-well plates (Greiner Cat. No. 784075) in 2 μL volumes, followed by 18 μL of assay mixture. Plates were incubated for 1 h at room temperature.

TR-FRET signals were measured on a Tecan Infinite M1000 Pro plate reader (excitation: 340 nm; emissions: 620 nm and 665 nm; 100 flashes/well; 500 μs integration time; 60 μs lag time). FRET was quantified as the 665/620 nm emission ratio × 100. Controls included 2% DMSO-treated wells and no-protein background wells (n = 3 per condition). For compounds that did not achieve ≥50% maximal inhibition, EC50 values are reported as relative EC50 values, calculated relative to the observed maximal inhibition (Imax).

### 4.6. Molecular Dynamics Simulations

All molecular dynamics simulations were performed using GROMACS 2025.3 with the CHARMM36m force field. System preparation, including topology generation, solvation, and ion placement, was performed using CHARMM-GUI. For unbiased binding site identification, six copies of NS-04 were placed around the SLIT2 LRR2 domain to maximize the probability of observing ligand-protein contacts. For pose stability assessment, the protein-ligand complex was prepared using the top-ranked pose from induced fit docking. All systems were solvated in an explicit TIP3P water box with 10 Å padding from the solute and neutralized to 150 mM NaCl using distance-based ion placement.

Following energy minimization, systems underwent equilibration under NVT and NPT ensembles according to CHARMM-GUI default protocols. Temperature was maintained at 300 K using the V-rescale thermostat, and pressure coupling during NPT equilibration employed the Parrinello-Rahman barostat. Covalent bonds involving hydrogen atoms were constrained using the LINCS algorithm, and long-range electrostatics were treated with particle mesh Ewald (PME). A 2-fs integration timestep was used for production runs under the Verlet cutoff scheme. Triplicate 100-ns production simulations were performed for both the multi-ligand binding site identification runs and the docked pose stability assessment. Ligand-protein contacts were quantified across all trajectories, and residues exhibiting sustained contacts were identified and visualized.

### 4.7. Induced Fit Docking

Induced fit docking (IFD) was performed using Schrödinger Maestro. The standard IFD protocol was employed, generating up to 20 poses per ligand. Ring conformations were sampled with an energy window of 2.5 kcal/mol. Initial Glide docking was performed with receptor and ligand van der Waals scaling factors of 0.50, with a maximum of 20 poses retained. Prime refinement was applied to residues within 5 Å of each ligand pose, with side chain optimization enabled. Glide redocking was performed into structures within 30 kcal/mol of the best structure and within the top 20 structures overall, using Standard Precision (SP) mode.

### 4.8. Chemistry

All reactions were performed under protection of N_2_ in oven-dried glassware unless water was applied as solvent. For Suzuki reactions, solvents are required to be degassed: a continuous flow of N_2_ was bubbled through the solvent for 3 hours, and a N_2_-filled balloon was attached to maintain an inert atmosphere during solvent withdrawal. The degassed solvent can be stored under nitrogen for extended periods without the need for further purging. Chromatographic purification was performed as flash chromatography with Combi-Flash^®^ Rf+ UV-VIS MS COMP using RediSep^®^ Silver normal phase silica gel columns and solvents indicated as eluent with default pressure. Fractions were collected based on UV absorption at 254 nm and/or 280 nm. Analytical thin layer chromatography (TLC) was performed on Whatman^®^ TLC silica gel UV254 (250 µm) TLC aluminum plates. Visualization was usually accomplished with UV light (254 nm and 280 nm). For the methyl amide product **6**, purifications were performed on XBridge BEH C18 OBD Prep Column 130Å, 5 μm, 19 mm x 150 mm on Waters^®^ HPLC-MS instruments with UV-VIS detector. HPLC was performed using a binary mobile phase consisting of water (A) and acetonitrile (B), each containing 0.1% formic acid. Proton and carbon nuclear magnetic resonance spectra (^1^H NMR and ^13^C NMR) were recorded on a cryo Bruker 500 MHz spectrometer or room temperature Bruker 600 MHz with solvent resonances as the internal standard (^1^H NMR: DMSO-d6 at 2.50 ppm. ^13^C NMR: DMSO-d6 at 39.52 ppm). 1H NMR data are reported as follows: chemical shift (ppm), multiplicity (s = singlet, d = doublet, dd = doublet of doublets, dt = doublet of triplets, ddd = doublet of doublet of doublets, t = triplet, m = multiplet, br = broad), coupling constants (Hz), and integration. Occasionally, tiny amount of solvents or additives of Prep-HPLC such as DCM, diethyl ether, methanol, acetonitrile, formic acid or acetone residues were also observed in the NMR spectrum. Mass spectra were obtained through ESI on Micromass Waters LCT Premier XE. The accurate mass analyses run in ESI mode were at a mass resolution of 10,000 to 15,000 FWHM and were calibrated using Leucine Enkephalin acetate as an internal standard.

#### Preparation of ethyl benzo[*d*]imidazole carboxylate intermediate (4)

General Step **a**: To a 4 mL vial charged with a stir bar was added to ethyl 4-fluoro-3-nitrobenzoate **1** (66.5 mg, 0.312 mmol, 1 eq.), amine or its HCl salt (0.328 mmol, 1.05 eq.), and Cs_2_CO_3_ (229 mg, 0.702 mmol, 2.25 eq.), followed by adding 1,4-dioxane (2 mL) and water (100 μL). The mixture was stirred vigorously and heated to 70°C for 1 hour and poured into 50 mL water, acidified to pH ∼7 and extracted with 25% THF in EtOAc (50 mL*2). The combined organic layer was washed with brine, dried over anhydrous Na_2_SO_4_, filtered and concentrated. The crude products **2** were moved forward for next step without further purification due to only one major consolidate spot was observed on TLC.

General Step **b**: To a 100 mL round bottom flask charged with a stir bar and all crude product **2** from step **a** was added 4 mL THF to completely dissolve all solid, followed by adding 30 mL EtOH. After the solution was purged with argon, palladium on carbon (10%) was added to the flask followed by vacuum and back-fill with pure hydrogen gas. The mixture was kept at room temperature for 3 hours to consume all starting material **2**. The mixture was then filtered and concentrated to produce crude product **3**, which could be moved forward for next step without further purification.

General Step **c**: To a 4 mL vial charged with a stir bar was added to all crude product **3** from step **b**, aryl aldehyde (0.328 mmol, 1.05 eq.), DMF (2 mL) and water (67 μL). The mixture was stirred vigorously for 5 mins, followed by adding Oxone^®^, monopersulfate compound (105 mg, 0.343 mmol, 1.1 eq.) and kept stirring for additional 40 mins. The reaction mixture was poured to 0.03M K_2_CO_3_ (50 mL), extracted with 25% THF in EtOAc (50 mL*3). The combined organic layer was washed with brine, dried over anhydrous Na_2_SO_4_, filtered and concentrated under reduced pressure to obtain crude product, which was absorbed onto a plug of silica gel and purified by chromatography (MeOH in DCM from 0%-5%).

#### Preparation of benzo[*d*]imidazole carboxylic acid (5)

General Step **d**: To a 20 mL vial charged with a stir bar was added to purified product **4** (0.12 mmol, 1 eq.), THF (3 mL), water (1 mL) and LiOH·H_2_O (20 mg, 0.48 mmol, 4 eq.). The mixture was stirred vigorously and heated to 60°C for overnight, after which was naturalized with 0.5 mL 1M HCl, poured into 50 mL water, exacted with 30% IPA in DCM (50 mL*3). The combined organic layer was washed with brine, dried over Na_2_SO_4_, filtered, and concentrated. The resulting solid was collected and triturated in 10% DCM in hexanes (5 mL) and 10% acetone in Et_2_O (5 mL) to product the pure product **5** without further purification.

#### Preparation of *N*-methyl benzo[*d*]imidazole carboxamide (6)

General Step **e**: To a 4 mL vial charged with a stir bar was added to product **5** (70 mmol, 1 eq.), NMP (1 mL), methylamine HCl salt (16.3 mg, 0.241 mmol, 3 eq.) and DIEA (60 mL, 0.345 mmol, 5 eq.). The mixture was stirred at room temperature for 5 minutes, followed by adding HATU (40 mg, 0.105 mmol, 1.5 eq.) and the kept stirred for overnight. 0.5 mL water (with 5% formic acid) was added into the mixture to quench the reaction. The mixture was purified under HPLC with the gradient of 5%-70% B, then the fractions with correct mass were collected and lyophilized to produce the final pure product **6**.

**Ethyl 2-(6-(1H-pyrrolo[2,3-b]pyridin-4-yl)quinolin-3-yl)-1-(2-(benzo[b]thiophen-3-yl)ethyl)-1H-benzo[d]imidazole-5-carboxylate (4a): 4a** was obtained as yellow solid (85 mg, 0.143 mmol, 46% over three steps) by applying general steps **a** to **c**, with **1**, 2-(benzo[b]thiophen-3-yl)ethan-1-amine HCl salt (70 mg, 0.328 mmol, 1.05 eq.) and 6-(1*H*-pyrrolo[2,3-b]pyridin-4-yl)quinoline-3-carbaldehyde (90 mg, 0.328 mmol, 1.05 eq.). **^1^H NMR** (500 MHz, DMSO) δ 11.92 (s, 1H), 8.93 (d, *J* = 2.2 Hz, 1H), 8.41 (d, *J* = 4.2 Hz, 2H), 8.36 (d, *J* = 1.6 Hz, 1H), 8.26 (d, *J* = 2.1 Hz, 1H), 8.21 (dd, *J* = 8.7, 2.0 Hz, 1H), 8.13 (d, *J* = 8.7 Hz, 1H), 8.01 (dd, *J* = 8.5, 1.6 Hz, 1H), 7.94 (d, *J* = 8.6 Hz, 1H), 7.67 (t, *J* = 3.0 Hz, 1H), 7.61 (d, *J* = 7.9 Hz, 1H), 7.38 (d, *J* = 4.9 Hz, 1H), 7.32 (d, *J* = 7.9 Hz, 1H), 6.95 – 6.89 (m, 2H), 6.89 – 6.84 (m, 1H), 6.81 (dd, *J* = 3.5, 1.9 Hz, 1H), 4.91 (t, *J* = 6.4 Hz, 2H), 4.38 (q, *J* = 7.1 Hz, 2H), 3.28 (t, *J* = 6.5 Hz, 2H), 1.39 (t, *J* = 7.1 Hz, 3H). **^13^C NMR** (126 MHz, DMSO) δ 166.10, 152.64, 149.71, 149.17, 146.76, 142.96, 142.35, 139.22, 139.11, 138.62, 137.69, 136.97, 136.00, 131.61, 130.60, 129.22, 127.98, 127.02, 126.31, 124.18, 124.15, 123.73, 123.70, 123.41, 122.90, 122.31, 120.94, 120.68, 117.31, 114.52, 111.23, 98.94, 60.52, 44.13, 27.87, 14.19. **LC-MS (ES+):** m/z = 594.35 [M+H]^+^

**2-(6-(1H-Pyrrolo[2,3-b]pyridin-4-yl)quinolin-3-yl)-1-(2-(benzo[b]thiophen-3-yl)ethyl)-1H-benzo[d]imidazole-5-carboxylic acid (5a): 5a** was obtained as matte yellow powder (45 mg, 80 mmol, 67%) by applying general step **d** with corresponding ethyl ester **4a** (71 mg, 0.12 mmol). **^1^H NMR** (500 MHz, DMSO) δ 12.83 (s, 1H), 11.92 (s, 1H), 8.94 (s, 1H), 8.41 (t, *J* = 4.0 Hz, 2H), 8.34 (d, *J* = 1.6 Hz, 1H), 8.26 (d, *J* = 2.1 Hz, 1H), 8.21 (dd, *J* = 8.7, 2.0 Hz, 1H), 8.13 (d, *J* = 8.7 Hz, 1H), 8.00 (dd, *J* = 8.4, 1.6 Hz, 1H), 7.92 (d, *J* = 8.5 Hz, 1H), 7.66 (t, *J* = 3.1 Hz, 1H), 7.61 (d, *J* = 7.9 Hz, 1H), 7.38 (d, *J* = 4.9 Hz, 1H), 7.32 (d, *J* = 7.9 Hz, 1H), 6.96 (s, 1H), 6.92 (t, *J* = 7.5 Hz, 1H), 6.87 (t, *J* = 7.5 Hz, 1H), 6.80 (dd, *J* = 3.6, 1.9 Hz, 1H), 4.90 (t, *J* = 6.5 Hz, 2H), 3.28 (t, *J* = 6.6 Hz, 2H). **^13^C NMR** (126 MHz, DMSO) δ 167.70, 152.42, 149.74, 146.74, 142.96, 142.34, 139.22, 139.11, 138.43, 137.70, 136.96, 135.99, 131.64, 130.57, 129.22, 127.97, 127.01, 126.32, 125.04, 124.15, 124.01, 123.69, 123.40, 122.98, 122.30, 121.08, 120.69, 117.31, 114.51, 111.01, 98.94, 44.09, 27.88. **HRMS (ESI)** m/z: [M+H]^+^ cal. for C34 H24 N5 O2 S, 566.1651; Found, 566.1638.

**Ethyl 1-(2-(benzo[b]thiophen-3-yl)ethyl)-2-(quinolin-3-yl)-1H-benzo[d]imidazole-5-carboxylate (4b): 4b** was obtained as yellow solid (73 mg, 0.153 mmol, 49% over three steps) by applying general steps **a** to **c**, with **1**, 2-(benzo[b]thiophen-3-yl)ethan-1-amine HCl salt (70 mg, 0.328 mmol, 1.05 eq.) and quinoline-3-carbaldehyde (52 mg, 0.328 mmol, 1.05 eq.). **^1^H NMR** (500 MHz, DMSO) δ 8.90 (d, *J* = 2.3 Hz, 1H), 8.36 – 8.32 (m, 1H), 8.29 (d, *J* = 2.3 Hz, 1H), 8.03 – 7.96 (m, 2H), 7.92 (d, *J* = 8.5 Hz, 1H), 7.89 – 7.80 (m, 2H), 7.67 (q, *J* = 7.9 Hz, 2H), 7.24 (d, *J* = 8.0 Hz, 1H), 7.00 (t, *J* = 7.6 Hz, 1H), 6.97 (s, 1H), 6.88 (t, *J* = 7.6 Hz, 1H), 4.84 (t, *J* = 6.6 Hz, 2H), 4.38 (q, *J* = 7.1 Hz, 2H), 3.27 (t, *J* = 6.7 Hz, 2H), 1.38 (t, *J* = 7.1 Hz, 3H). **^13^C NMR** (126 MHz, DMSO) δ 166.68, 153.29, 149.84, 147.75, 142.91, 139.84, 139.18, 138.33, 136.30, 132.20, 131.13, 129.16, 127.67, 126.76, 124.72, 124.67, 124.37, 124.26, 124.04, 123.11, 123.02, 121.49, 121.19, 111.78, 61.10, 44.82, 28.35, 14.76. **LC-MS (ES+):** m/z = 478.24 [M+H]^+^

**1-(2-(Benzo[b]thiophen-3-yl)ethyl)-2-(quinolin-3-yl)-1H-benzo[d]imidazole-5-carboxylic acid (5b): 5b** was obtained as yellow powder (41 mg, 91 mmol, 75%) by applying general step **d** with corresponding ethyl ester **4b** (58 mg, 0.12 mmol). **^1^H NMR** (500 MHz, DMSO) δ 12.82 (s, 1H), 8.91 (d, *J* = 2.2 Hz, 1H), 8.31 (dd, *J* = 9.8, 1.9 Hz, 2H), 8.03 – 7.95 (m, 2H), 7.92 – 7.78 (m, 3H), 7.68 (p, *J* = 7.4 Hz, 2H), 7.24 (d, *J* = 8.0 Hz, 1H), 7.00 (t, *J* = 9.6 Hz, 2H), 6.88 (t, *J* = 7.5 Hz, 1H), 4.83 (t, *J* = 6.6 Hz, 2H), 3.27 (t, *J* = 6.7 Hz, 2H). **^13^C NMR** (126 MHz, DMSO) δ 167.69, 152.49, 149.29, 147.14, 142.32, 139.26, 138.40, 137.75, 135.70, 131.65, 130.52, 128.57, 127.08, 126.19, 124.99, 124.10, 123.96, 123.78, 123.45, 122.60, 122.43, 121.05, 120.62, 110.98, 44.21, 27.78. **HRMS (ESI)** m/z: [M+H]^+^ cal. for C27 H20 N3 O2 S, 450.1276; Found, 450.1284.

**1-(2-(Benzo[b]thiophen-3-yl)ethyl)-N-methyl-2-(quinolin-3-yl)-1H-benzo[d]imidazole-5-carboxamide (6b): 6b** was obtained as yellow solid (25 mg, 54 mmol, 76%) by applying general step **e** with corresponding acid **5b** (32 mg, 71 mmol). ^1^H NMR (500 MHz, DMSO) δ 8.91 (d, *J* = 2.2 Hz, 1H), 8.50 (q, *J* = 4.4 Hz, 1H), 8.28 (dd, *J* = 9.1, 1.9 Hz, 2H), 8.01 (d, *J* = 8.4 Hz, 1H), 7.91 (dd, *J* = 8.5, 1.6 Hz, 1H), 7.88 – 7.80 (m, 3H), 7.72 – 7.63 (m, 2H), 7.24 (d, *J* = 8.0 Hz, 1H), 7.05 – 6.97 (m, 2H), 6.89 (t, *J* = 7.5 Hz, 1H), 4.81 (t, *J* = 6.6 Hz, 2H), 3.27 (t, *J* = 6.7 Hz, 2H), 2.86 (d, *J* = 4.5 Hz, 3H). ^13^C NMR (126 MHz, DMSO) δ 166.89, 152.09, 149.45, 147.18, 142.38, 139.33, 137.85, 137.21, 135.66, 131.80, 130.54, 129.00, 128.64, 128.61, 127.13, 126.28, 124.10, 123.86, 123.53, 122.84, 122.52, 122.27, 120.72, 118.43, 110.74, 44.22, 27.88, 26.37. **HRMS (ESI)** m/z: [M+H]^+^ cal. for C28 H23 N4 O S, 463.1593; Found, 463.1588.

**Ethyl 2-(6-(1H-pyrrolo[2,3-b]pyridin-4-yl)quinolin-3-yl)-1-methyl-1H-benzo[d]imidazole-5-carboxylate (4c): 4c** was obtained as yellow solid (71 mg, 0.159 mmol, 51% over three steps) by applying general steps **a** to **c**, with methylamine HCl salt (42 mg, 0.624 mmol, 2 eq.) and 6-(1*H*-pyrrolo[2,3-b]pyridin-4-yl)quinoline-3-carbaldehyde (90 mg, 0.328 mmol, 1.05 eq.). **^1^H NMR** (500 MHz, DMSO) δ 11.91 (s, 1H), 9.46 (d, *J* = 2.2 Hz, 1H), 9.11 (d, *J* = 2.2 Hz, 1H), 8.63 (t, *J* = 1.4 Hz, 1H), 8.41 – 8.35 (m, 2H), 8.30 (d, *J* = 1.4 Hz, 2H), 7.99 (dd, *J* = 8.5, 1.7 Hz, 1H), 7.84 (d, *J* = 8.5 Hz, 1H), 7.64 (t, *J* = 3.0 Hz, 1H), 7.39 (d, *J* = 4.9 Hz, 1H), 6.82 (dd, *J* = 3.5, 1.8 Hz, 1H), 4.36 (q, *J* = 7.1 Hz, 2H), 4.10 (s, 3H), 1.38 (t, *J* = 7.1 Hz, 3H). **^13^C NMR** (126 MHz, DMSO) δ 166.10, 152.46, 150.65, 149.19, 147.08, 142.92, 142.03, 139.89, 138.95, 137.30, 136.97, 131.02, 129.42, 128.26, 127.02, 126.89, 124.04, 123.66, 123.22, 120.77, 117.27, 114.51, 110.88, 99.02, 60.51, 32.13, 14.16. **LC-MS (ES+):** m/z = 448.34 [M+H]^+^

**2-(6-(1H-Pyrrolo[2,3-b]pyridin-4-yl)quinolin-3-yl)-1-methyl-1H-benzo[d]imidazole-5-carboxylic acid (5c): 5c** was obtained as yellow powder (36 mg, 86 mmol, 71%) by applying general step **d** with corresponding ethyl ester **4c** (54 mg, 0.12 mmol). **^1^H NMR** (500 MHz, DMSO) δ 11.93 (s, 1H), 9.46 (s, 1H), 9.12 (s, 1H), 8.64 (s, 1H), 8.40 – 8.24 (m, 4H), 7.99 (d, *J* = 8.5 Hz, 1H), 7.78 (d, *J* = 8.5 Hz, 1H), 7.64 (t, *J* = 2.9 Hz, 1H), 7.40 (d, *J* = 4.9 Hz, 1H), 6.82 (d, *J* = 3.4 Hz, 1H), 4.10 (s, 3H). **^13^C NMR** (126 MHz, DMSO) δ 168.02, 152.01, 150.67, 149.17, 147.04, 142.91, 142.03, 139.38, 138.96, 137.27, 136.90, 130.95, 129.41, 128.24, 127.02, 126.92, 126.33, 124.05, 123.38, 120.78, 117.27, 114.50, 110.40, 99.00, 32.06. **HRMS (ESI)** m/z: [M+H]^+^ cal. for C25 H18 N5 O2, 420.1460; Found, 420.1442.

**2-(6-(1H-Pyrrolo[2,3-b]pyridin-4-yl)quinolin-3-yl)-N,1-dimethyl-1H-benzo[d]imidazole-5-carboxamide (6c): 6c** was obtained as yellow solid (24 mg, 55 mmol, 80%) by applying general step **e** with corresponding acid **5c** (29 mg, 69 mmol). **^1^H NMR** (500 MHz, DMSO) δ 11.91 (s, 1H), 9.45 (d, *J* = 2.3 Hz, 1H), 9.09 (d, *J* = 2.4 Hz, 1H), 8.62 (s, 1H), 8.49 (q, *J* = 4.4 Hz, 1H), 8.38 (d, *J* = 4.8 Hz, 1H), 8.29 (d, *J* = 2.5 Hz, 3H), 7.90 (dd, *J* = 8.5, 1.7 Hz, 1H), 7.77 (d, *J* = 8.5 Hz, 1H), 7.64 (t, *J* = 2.7 Hz, 1H), 7.39 (d, *J* = 4.9 Hz, 1H), 6.81 (d, *J* = 3.5 Hz, 1H), 4.08 (s, 3H), 2.84 (d, *J* = 4.4 Hz, 3H). **^13^C NMR** (126 MHz, DMSO) δ 166.91, 151.83, 150.79, 149.25, 147.10, 143.00, 142.09, 139.05, 138.55, 137.35, 136.86, 131.00, 129.49, 128.94, 128.30, 127.09, 127.00, 123.51, 122.26, 118.30, 117.35, 114.59, 110.43, 99.10, 32.08, 26.36. **HRMS (ESI)** m/z: [M+H]^+^ cal. for C26 H21 N6 O, 433.1777; Found 433.1763.

## Data Availability

Data will be made available on request.

## Acknowledgments

We acknowledge funding support by R01CA293456 (PI Gabr) from the National Cancer Institute (NCI).

