## Supporting Information for "Discovery and Optimization of Small Molecule Inhibitors of the SLIT2/ROBO1 Protein-Protein Interaction Using DNA-Encoded Libraries"

*Electronic Supplementary Information*

| **Contents** |  |
| --- | --- |
| Characterization data of procured compounds from AIphaMa | S2 |
| Characterization data of in-house synthesized compounds | S10 |

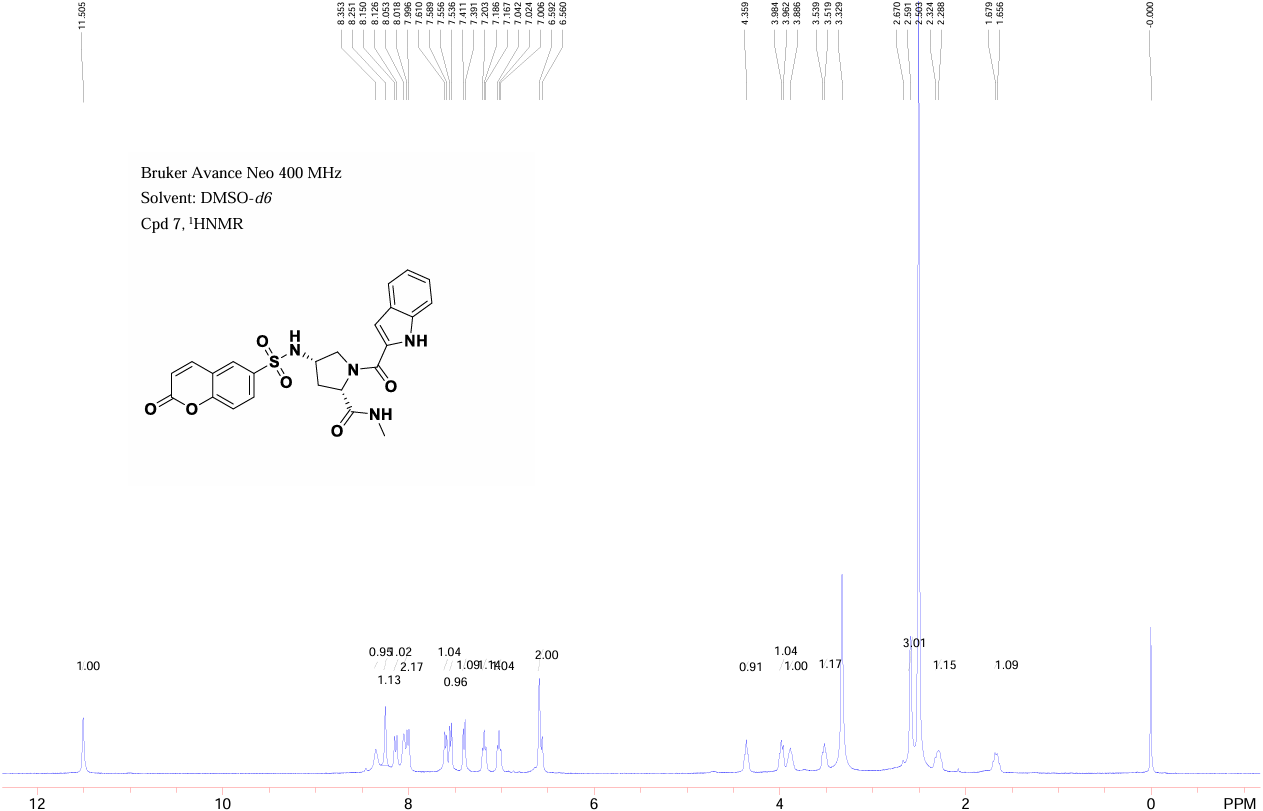
**NS-01**

**1H NMR**

**NS-01**

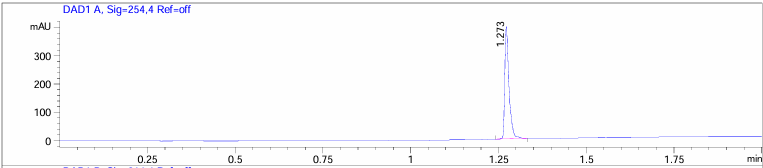

**NS-01**

**LC-MS**

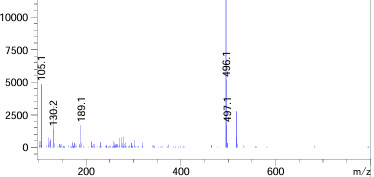

**NS-01**

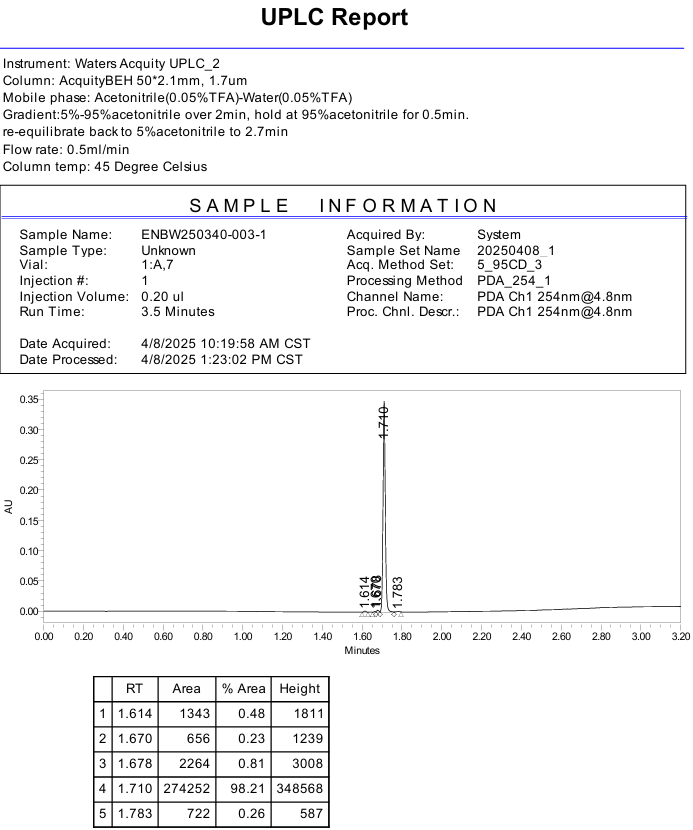

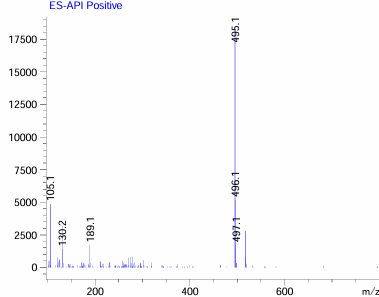

**NS-01**

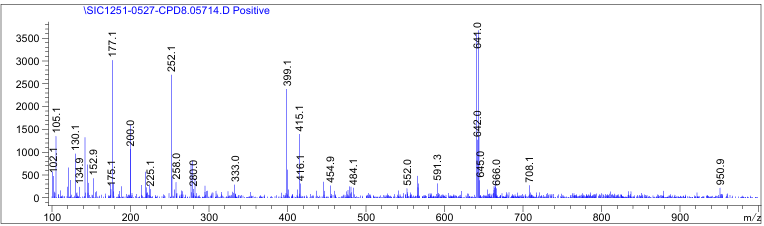

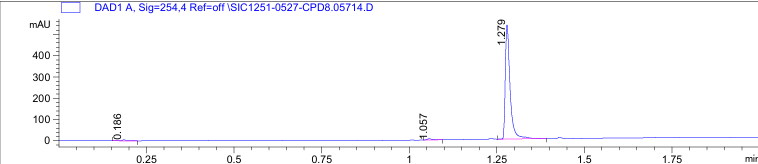

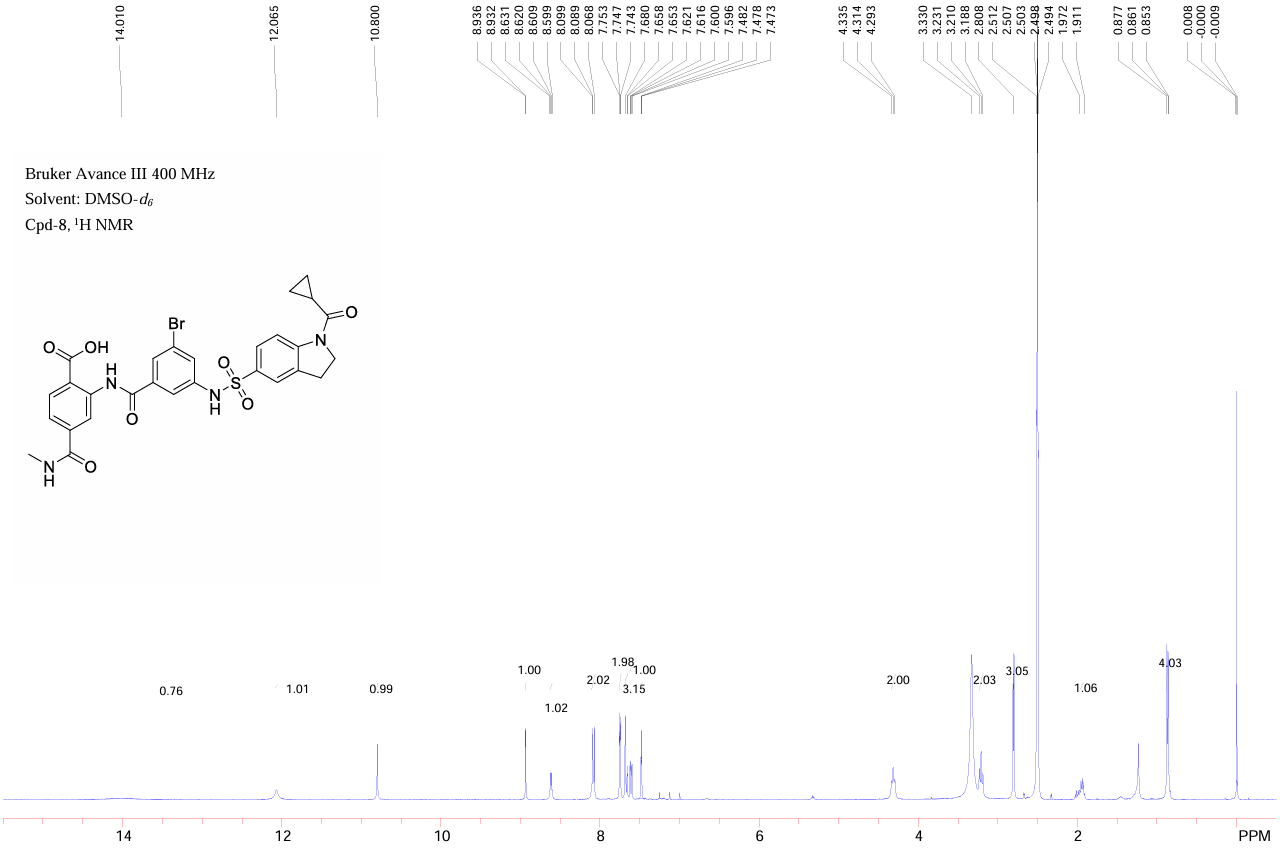
**NS-02**

**NS-02**

**NS-02**

**LC-MS**

**1H NMR**

**NS-02**

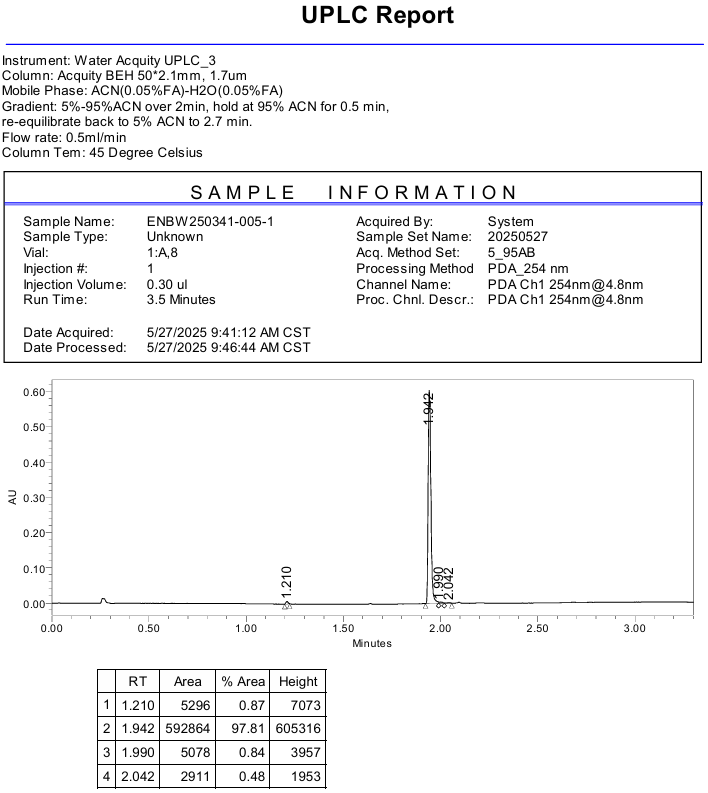

**NS-02**

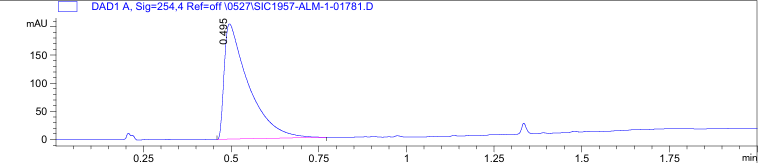

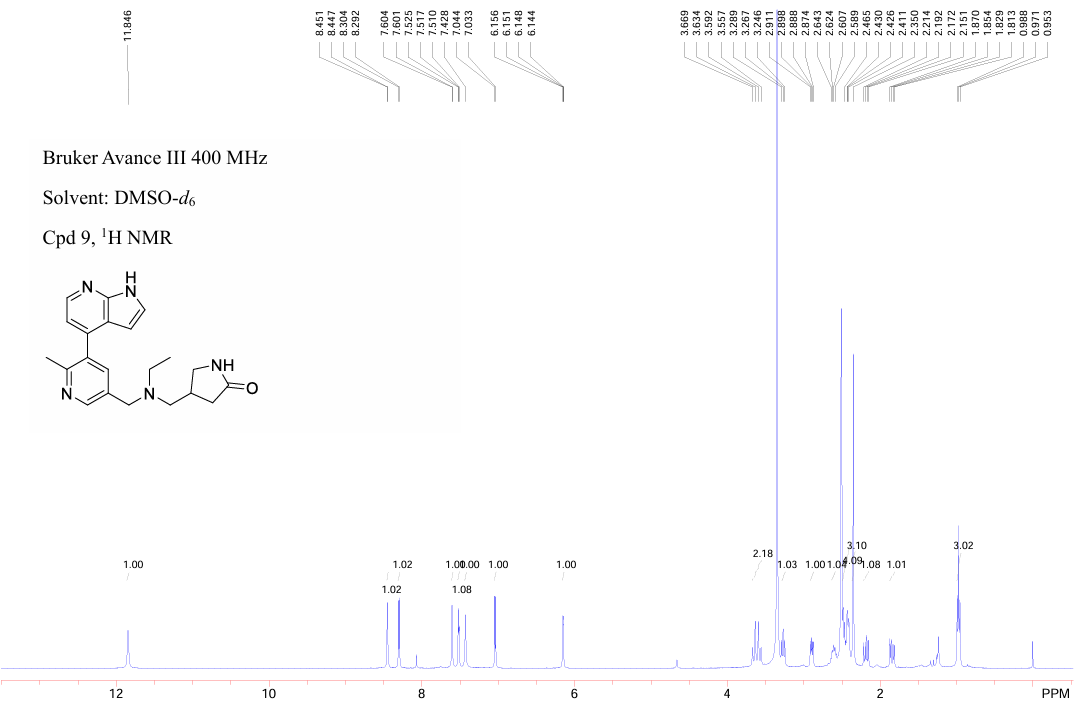
**NS-03**

**NS-03**

**LC-MS**

**1H NMR**

**NS-03**

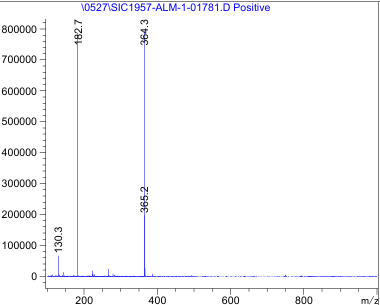

**NS-03**

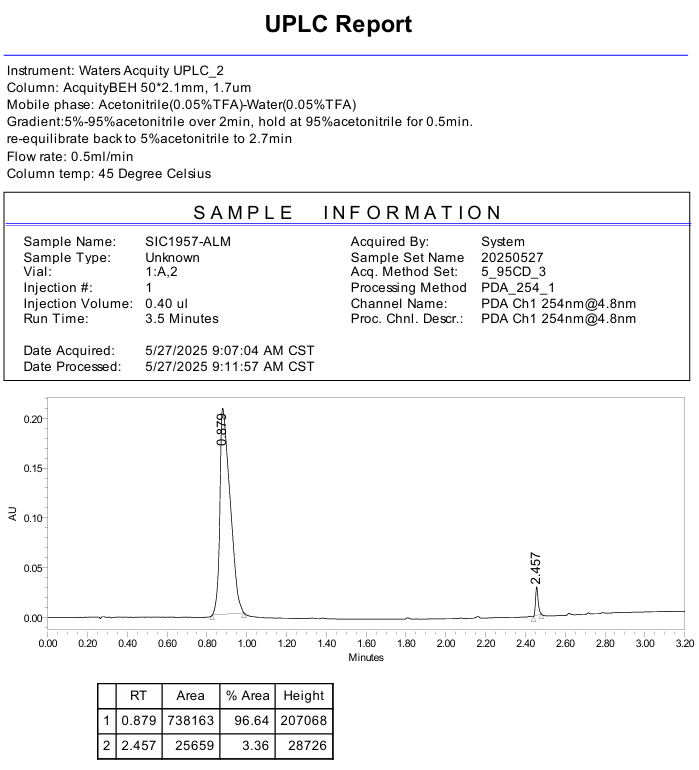

**NS-03**

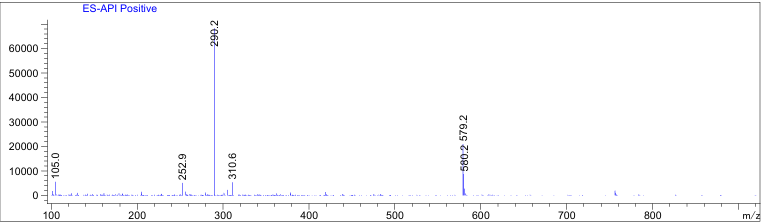

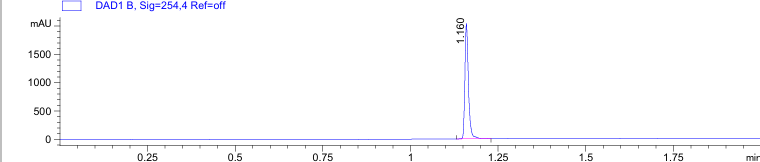

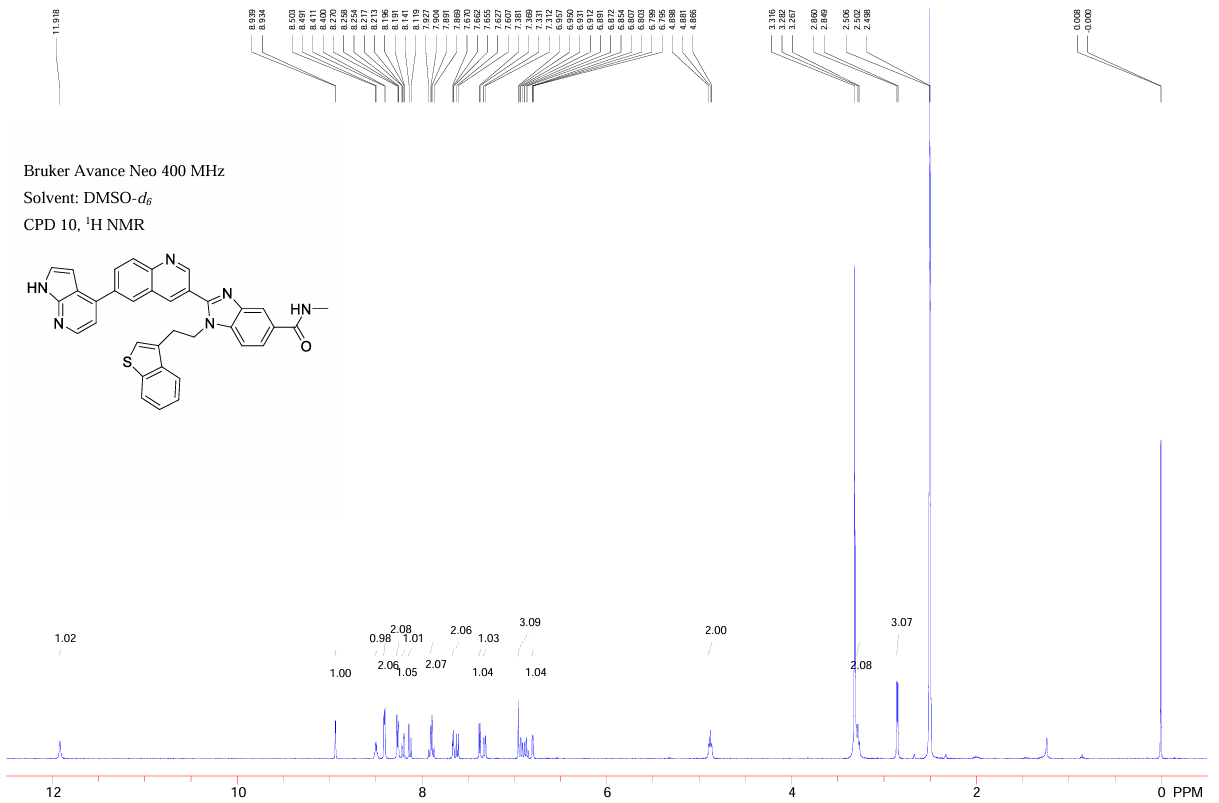
**NS-04**

**1H NMR**

**NS-04**

**LC-MS**

**NS-04**

**NS-04**

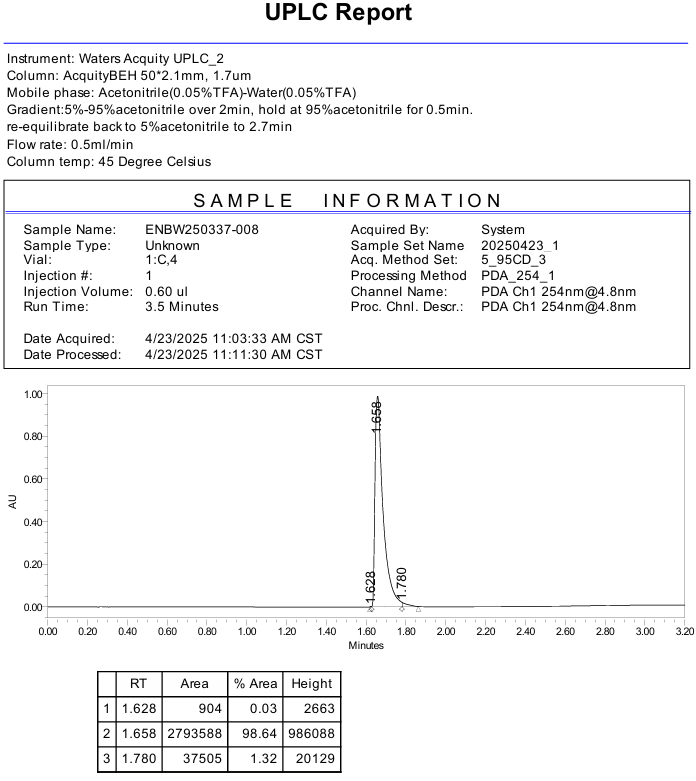

**NS-04**

**1H, 13C NMR of synthesized compounds**

**HRMS for compounds 5 and 6**

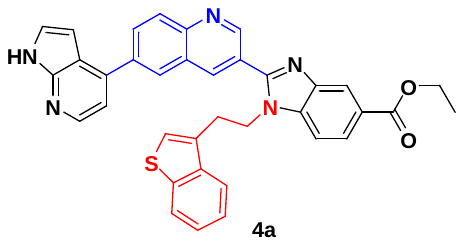

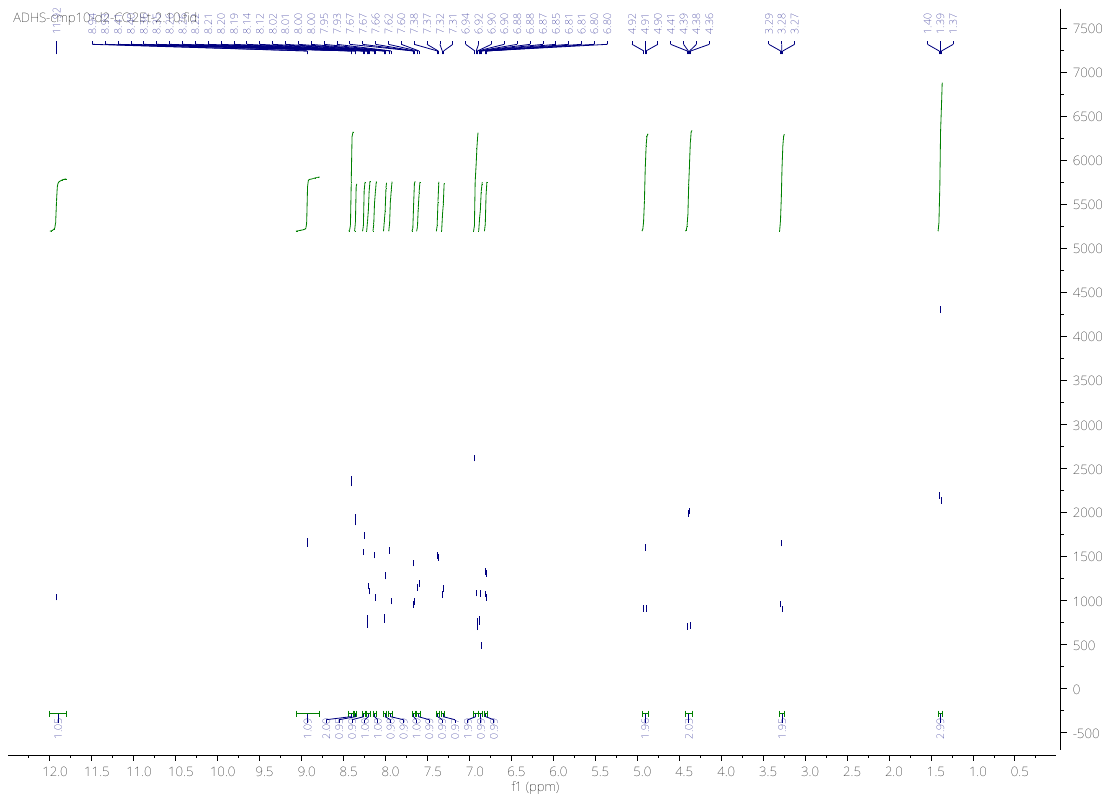

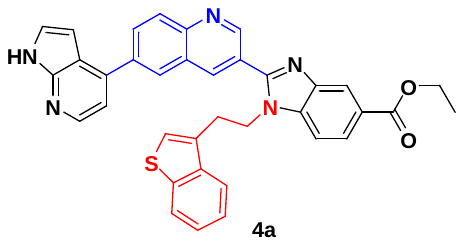

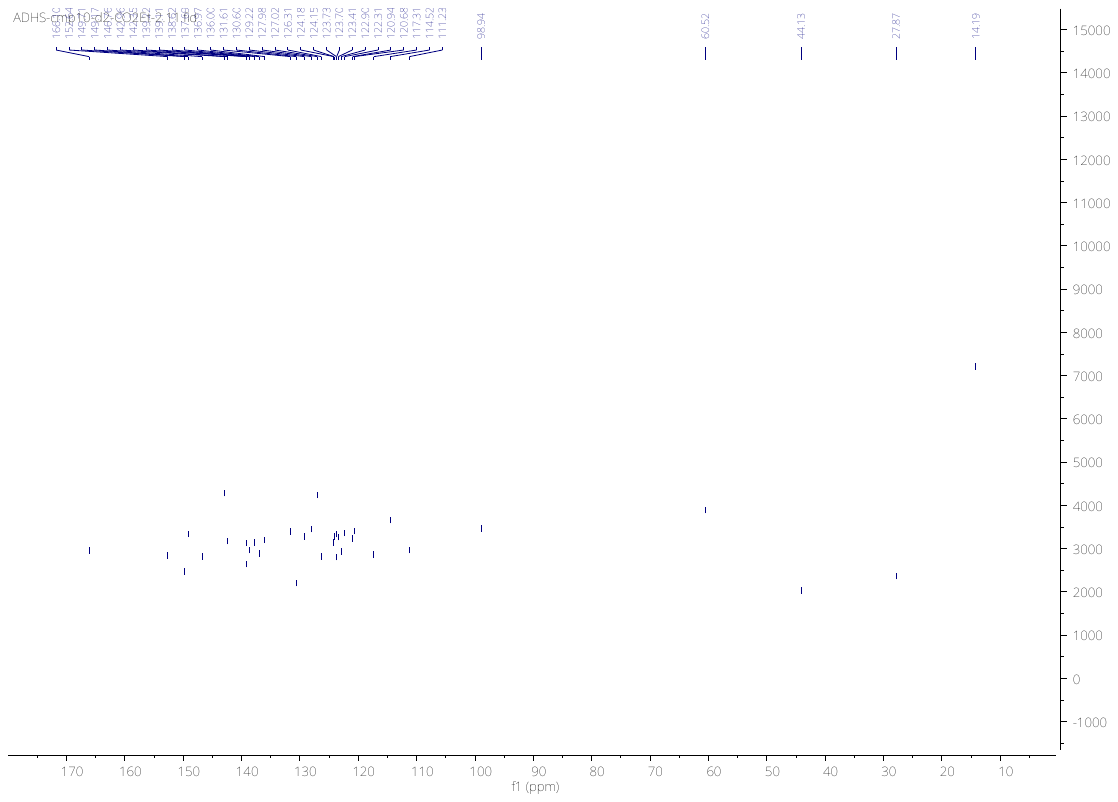

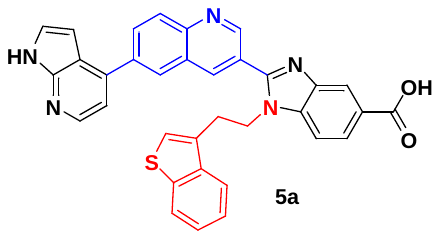

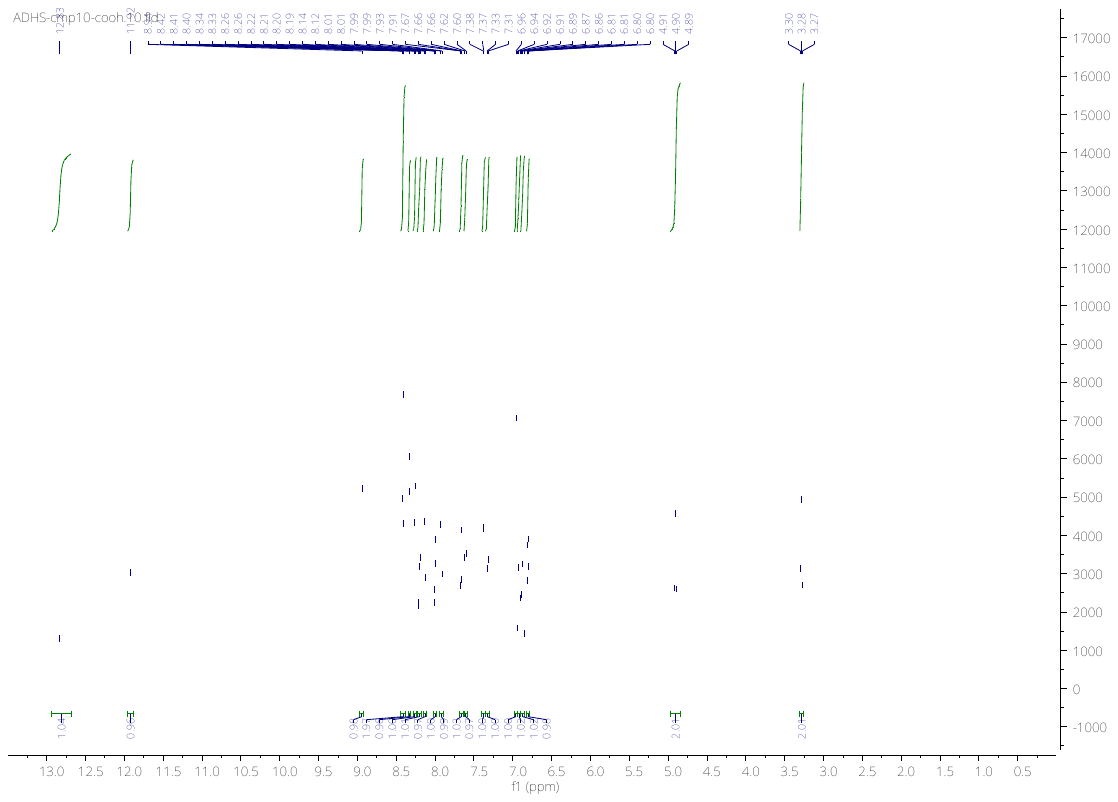

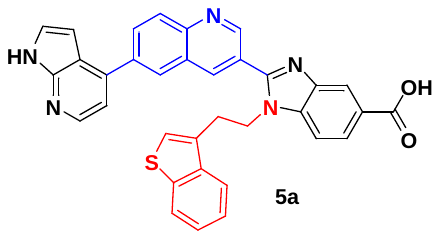

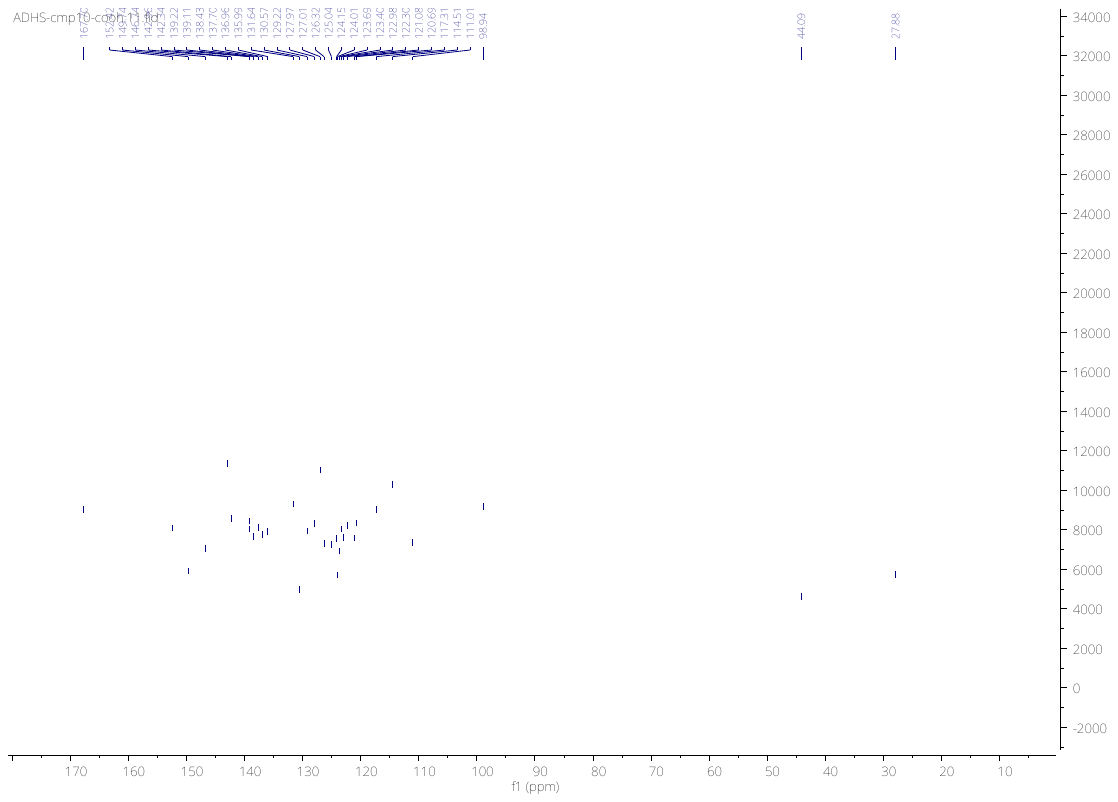

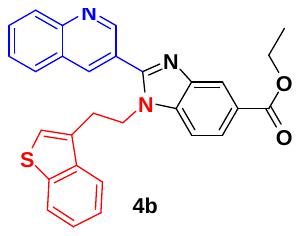

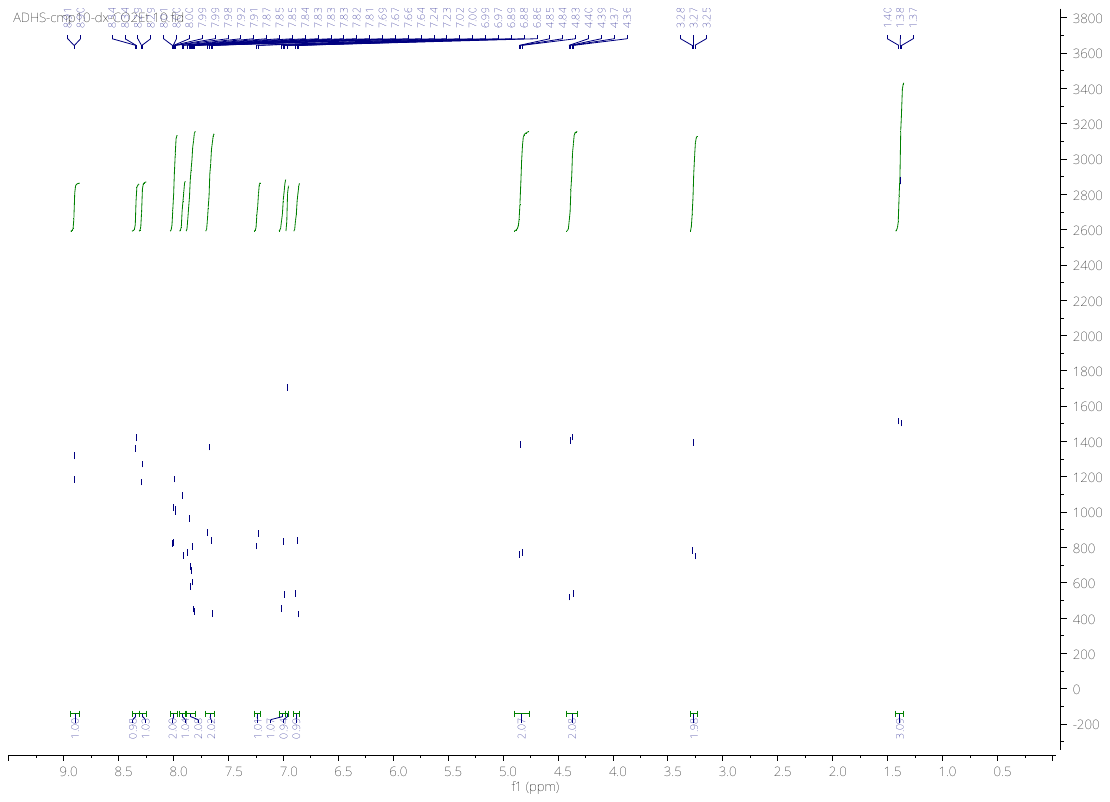

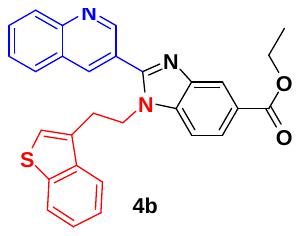

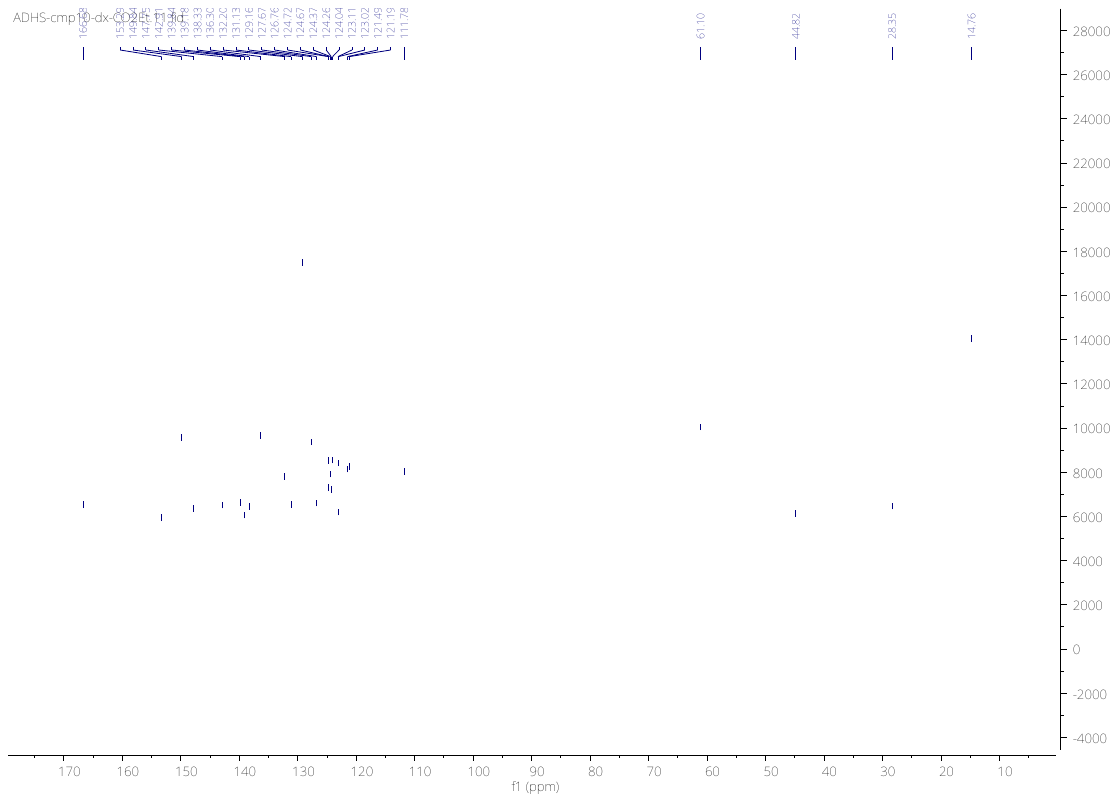

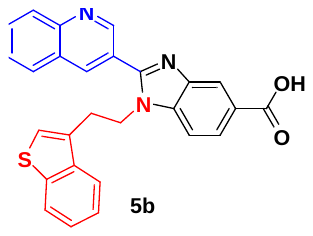

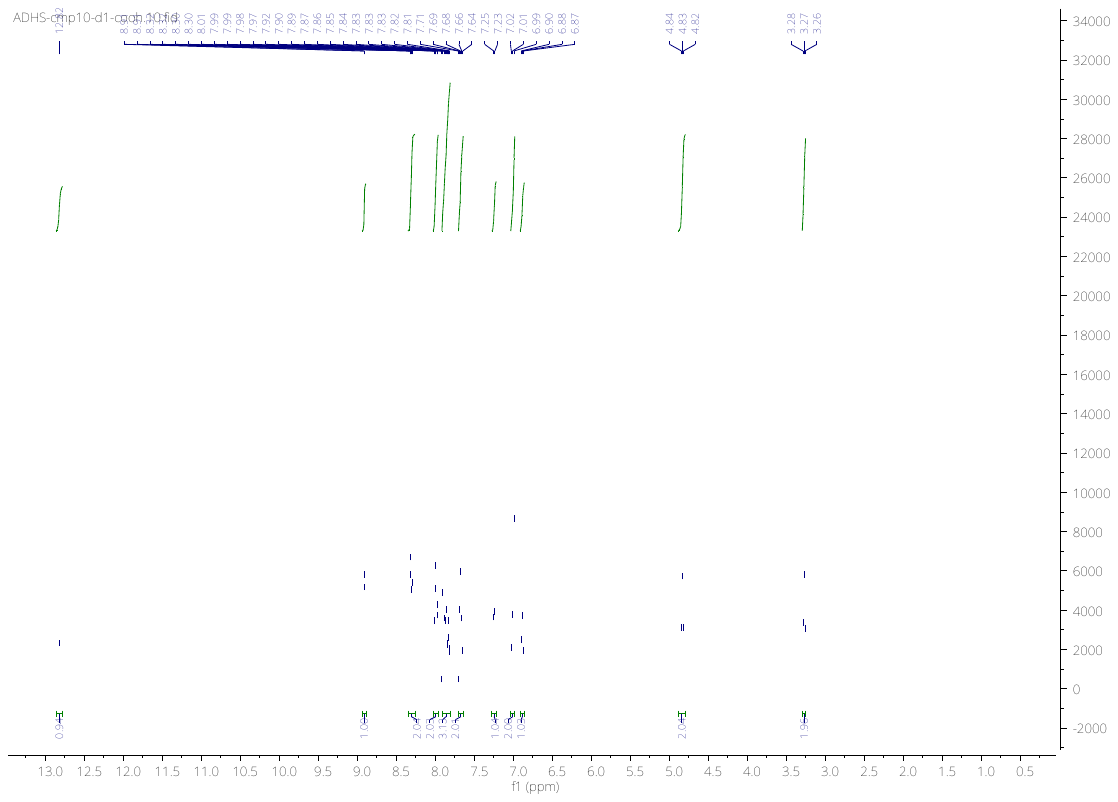

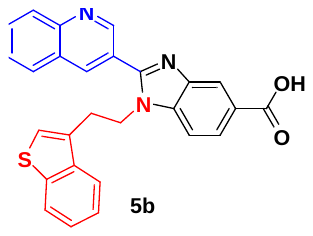

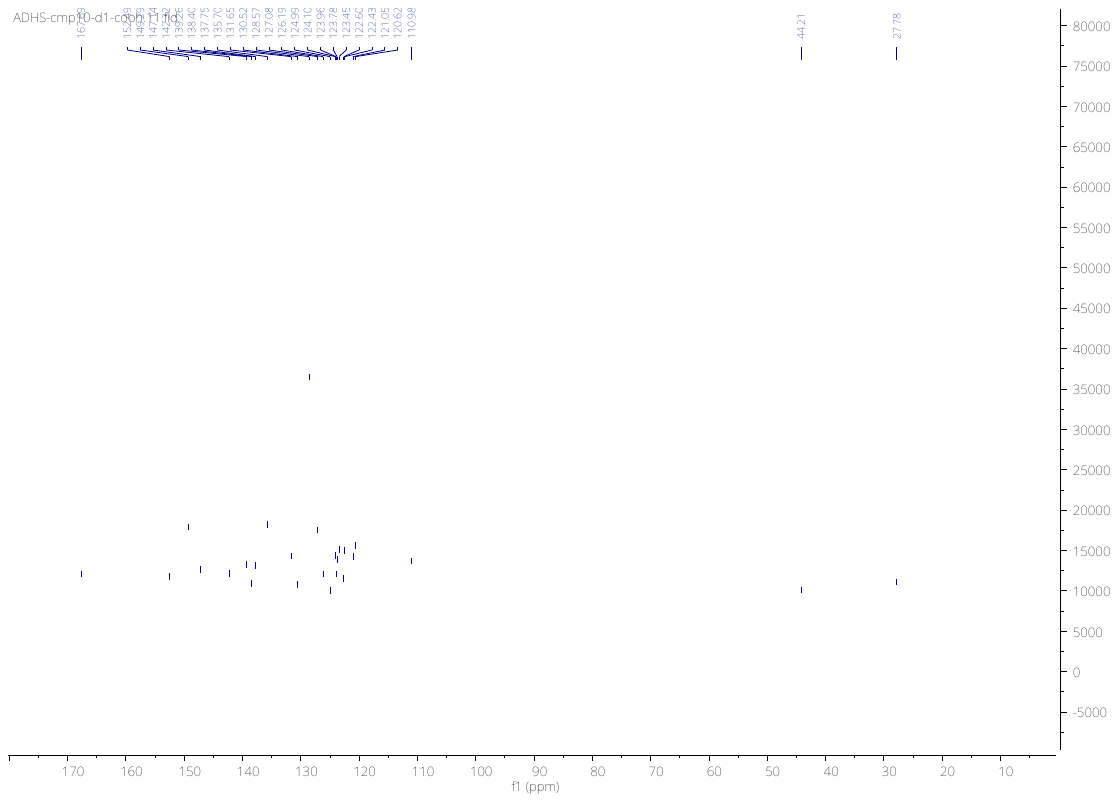

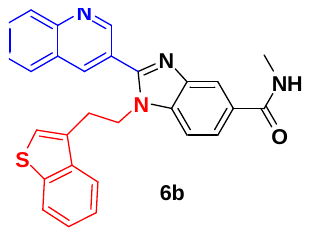

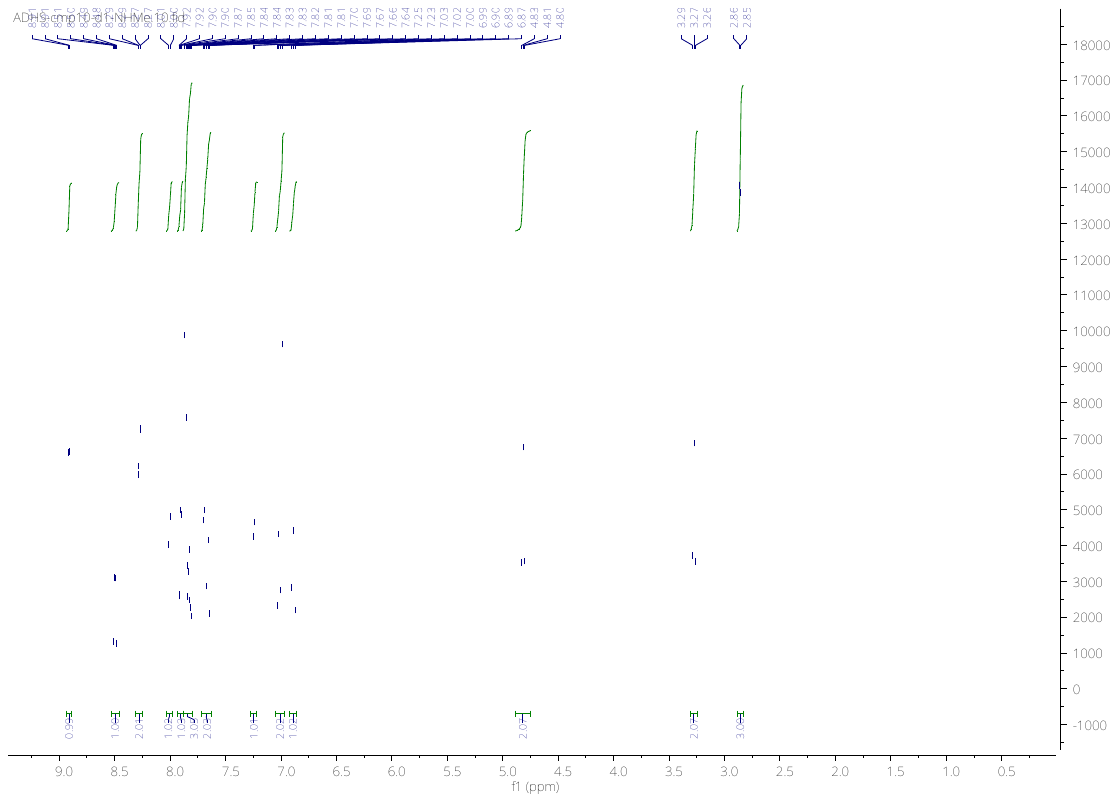

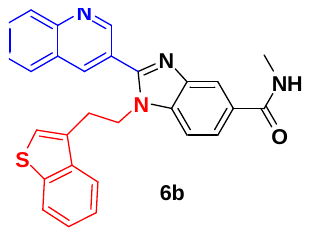

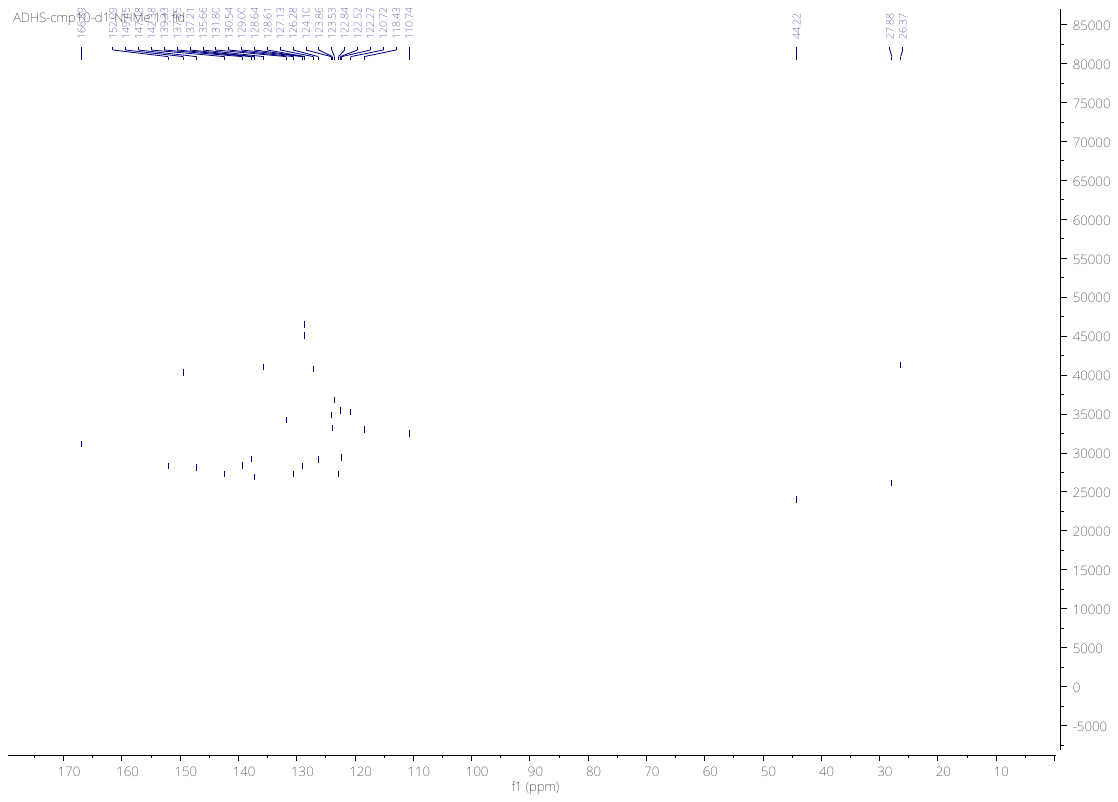

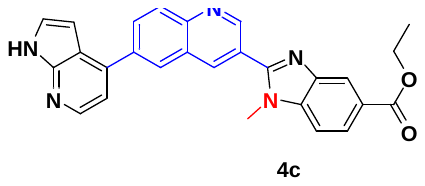

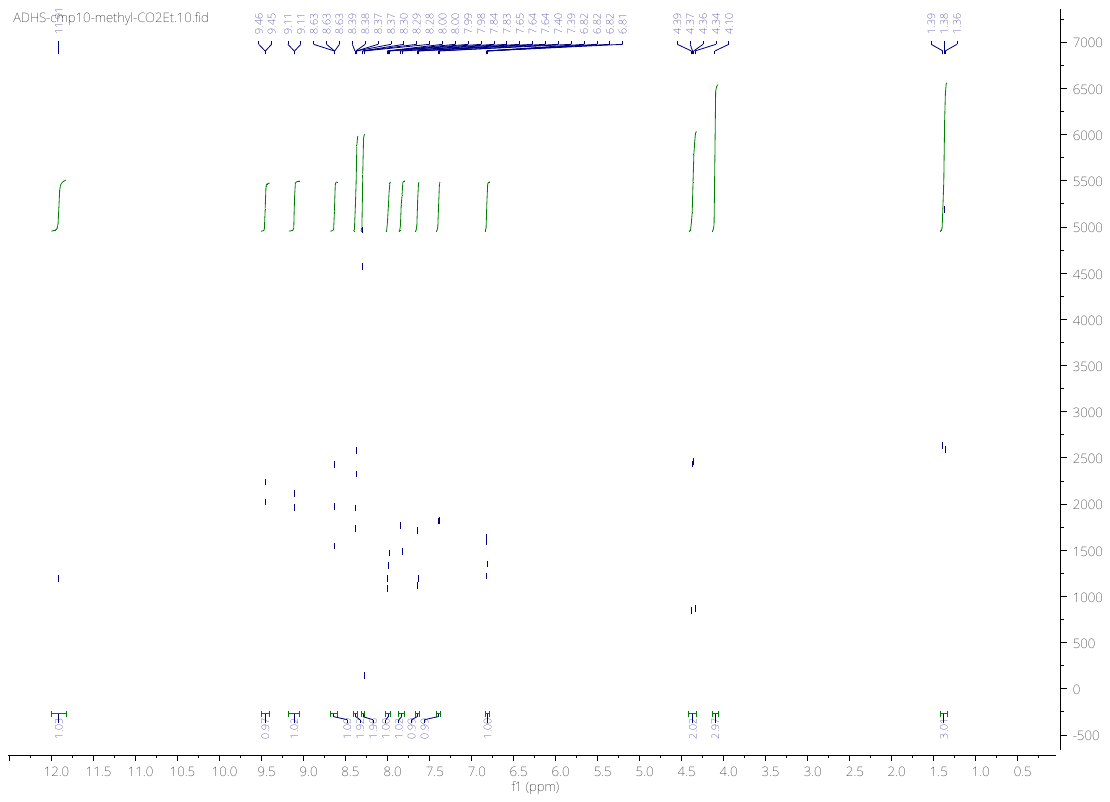

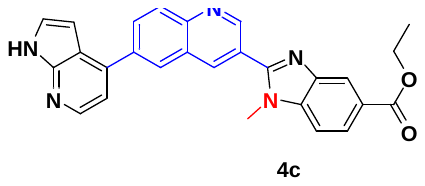

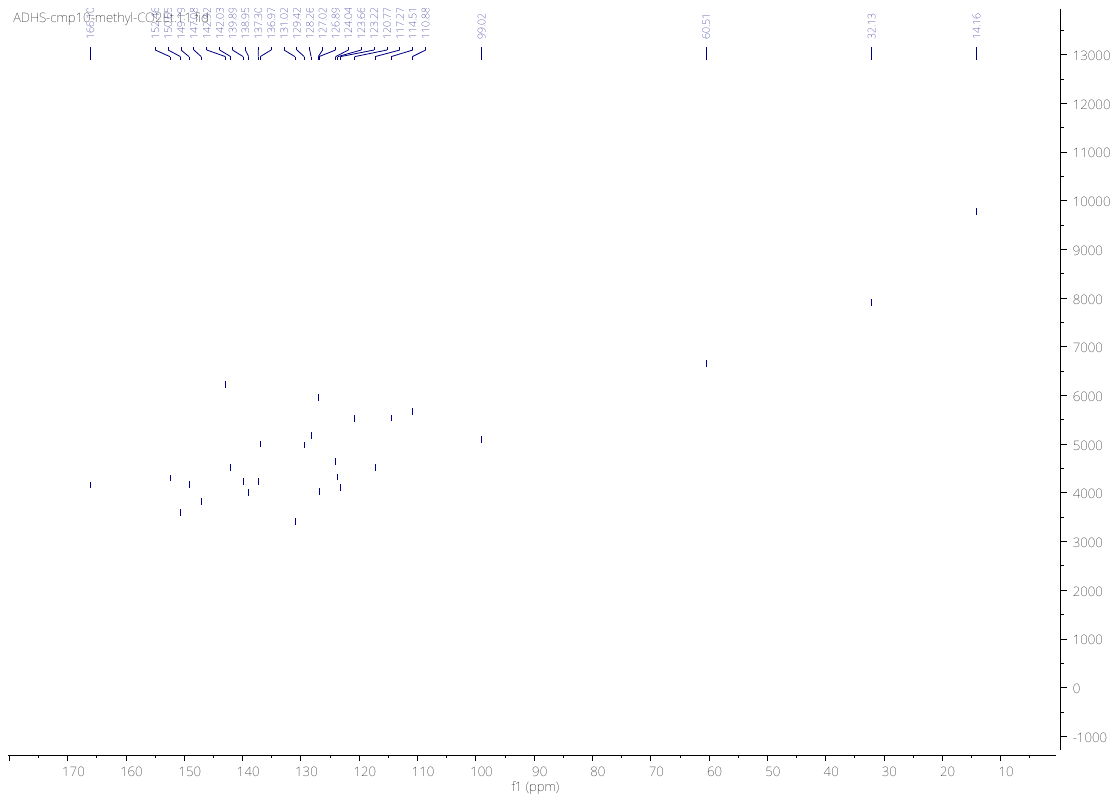

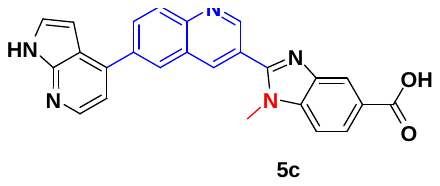

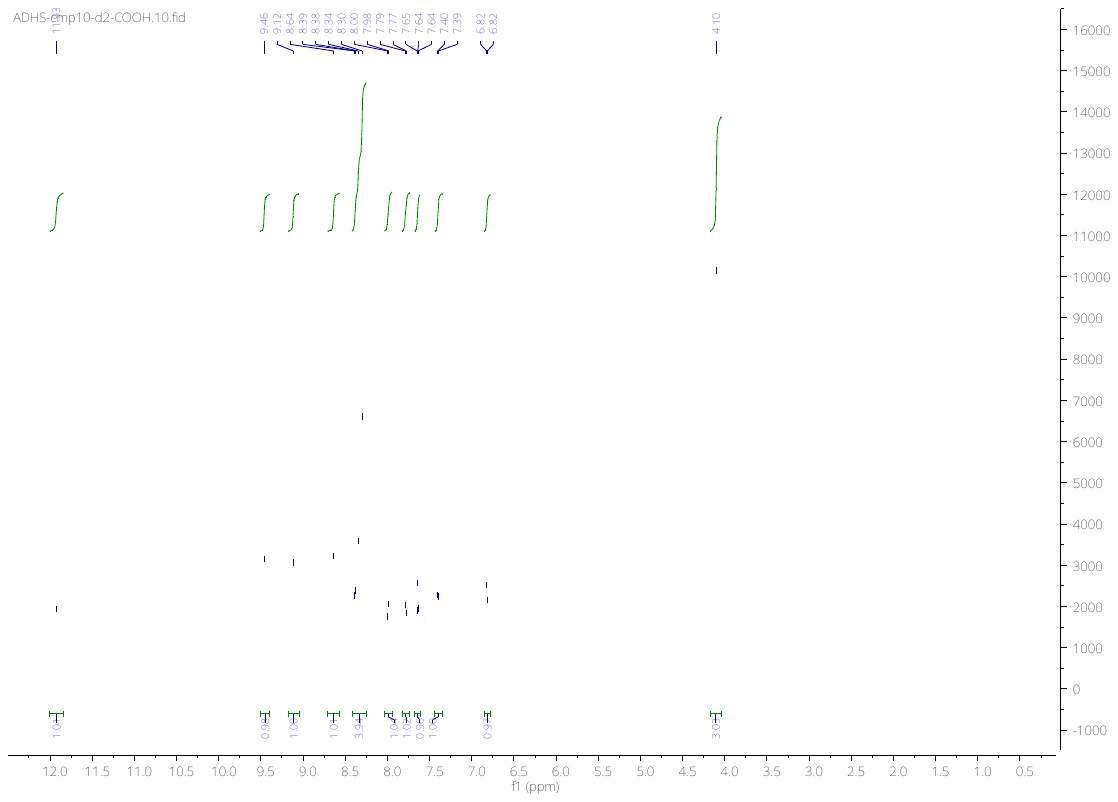

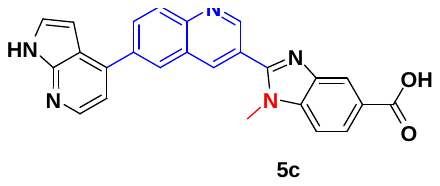

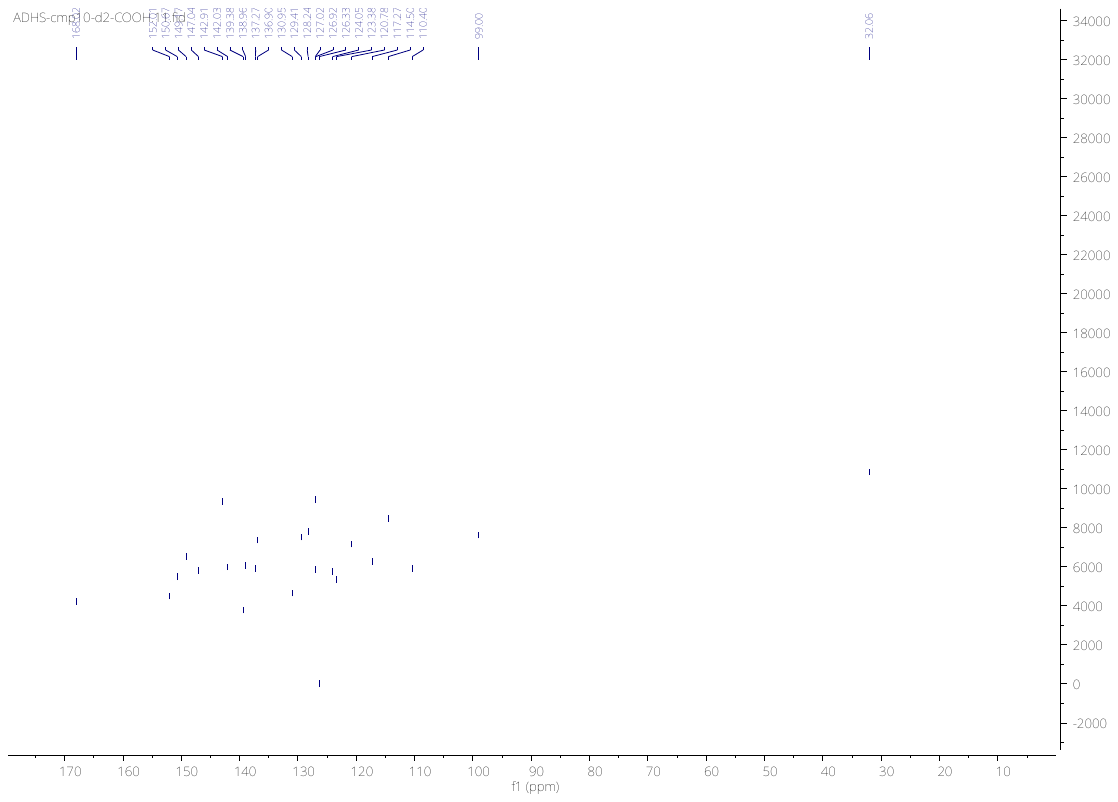

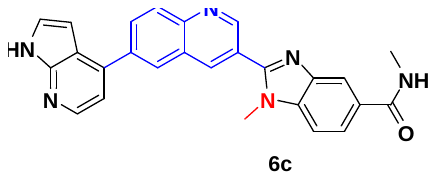

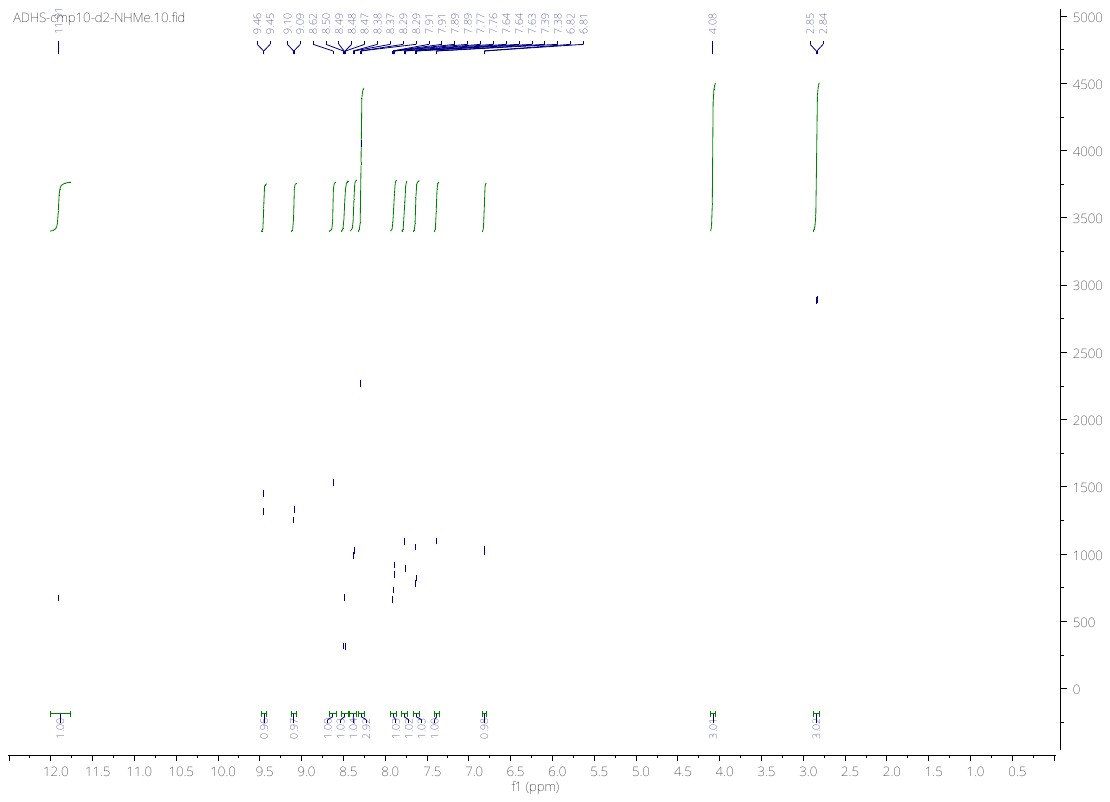

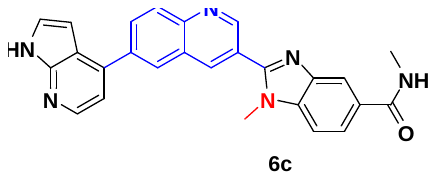

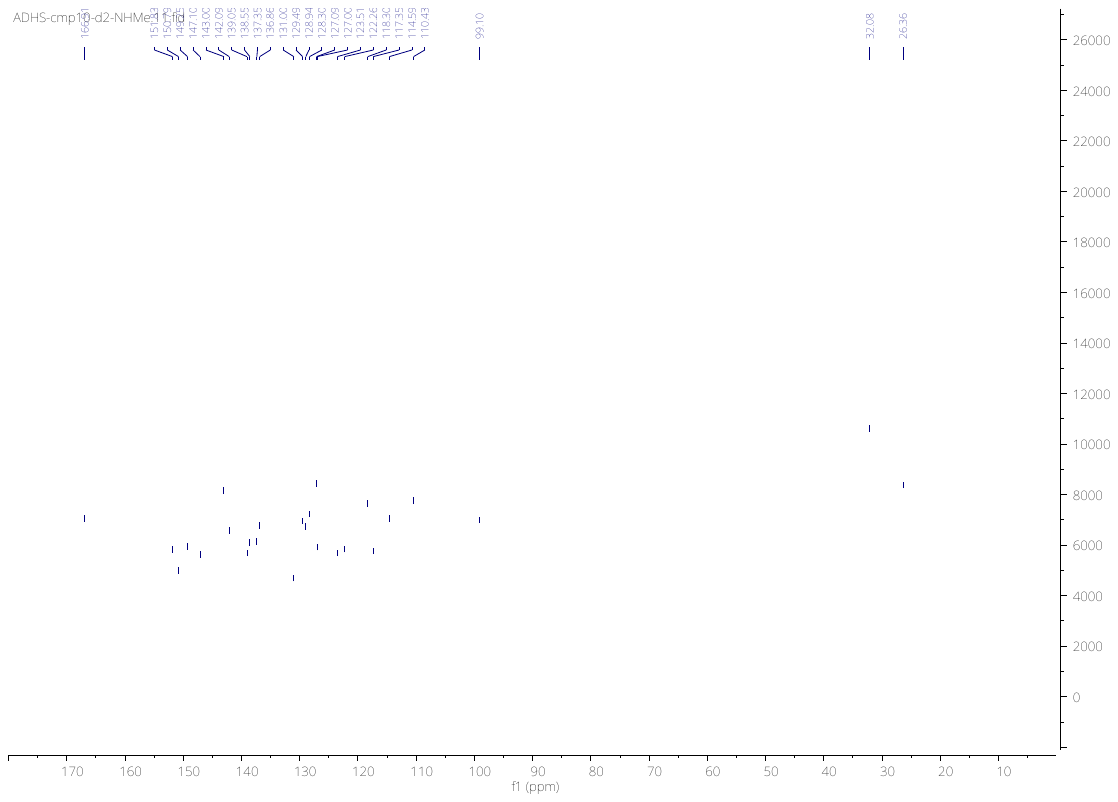

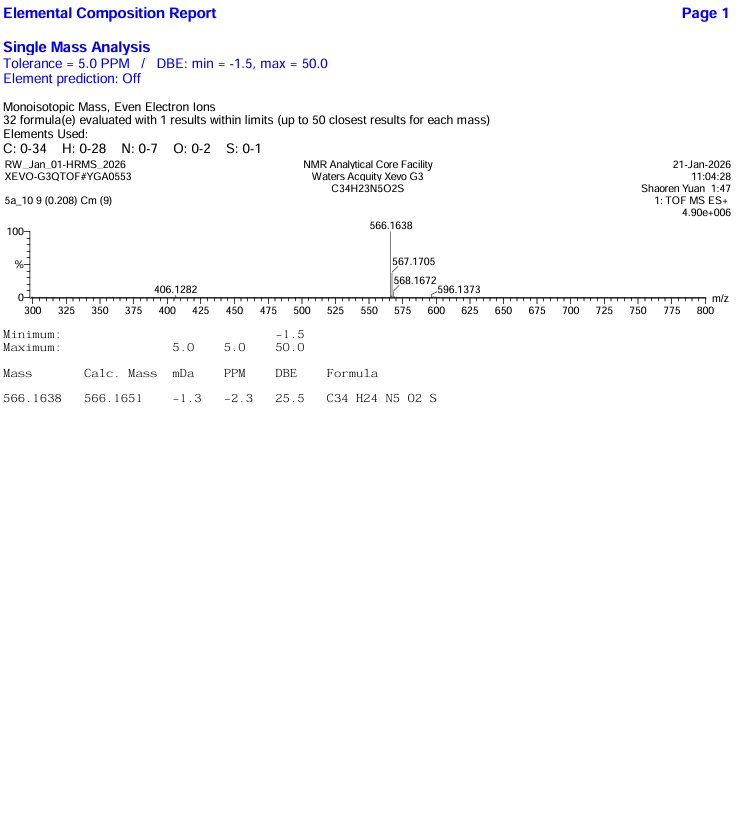

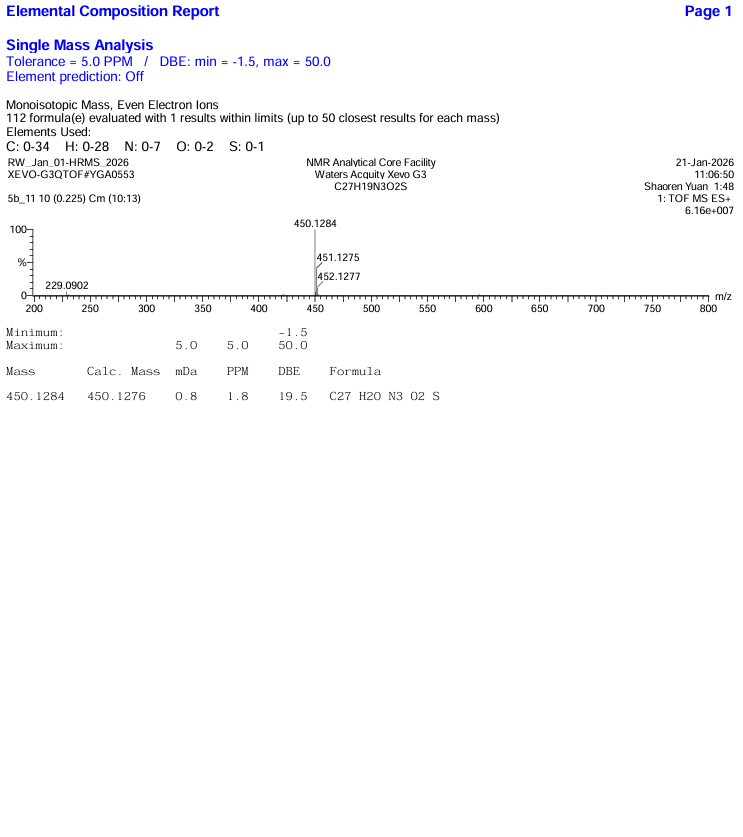

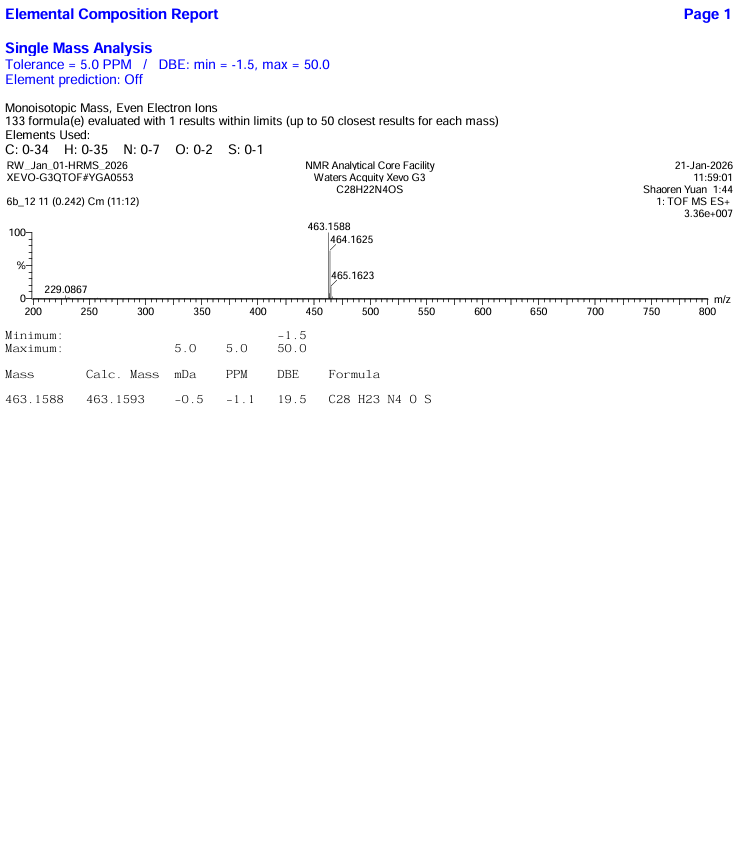

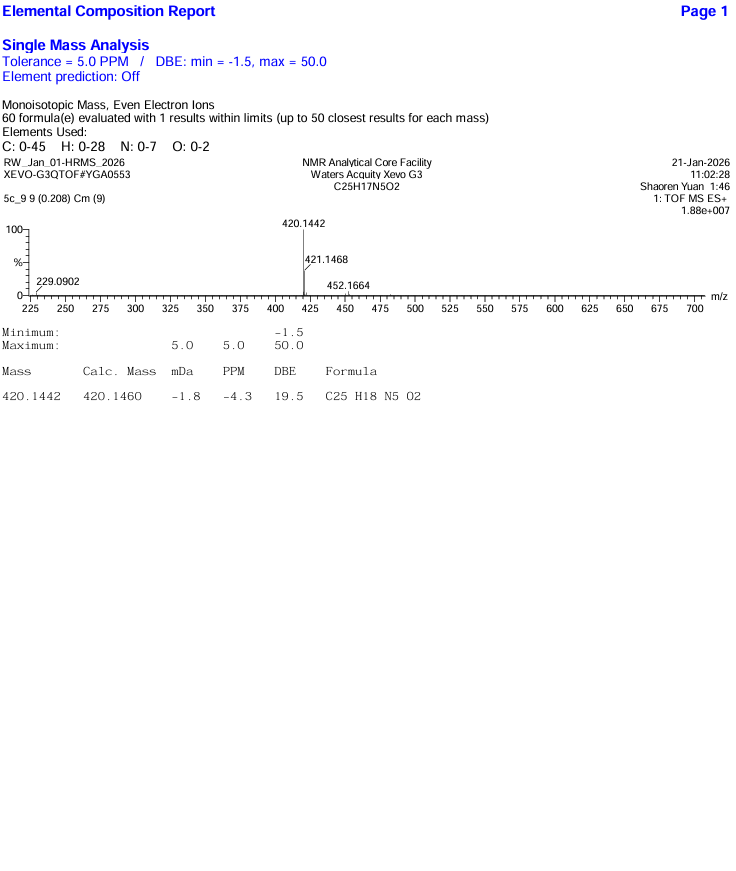

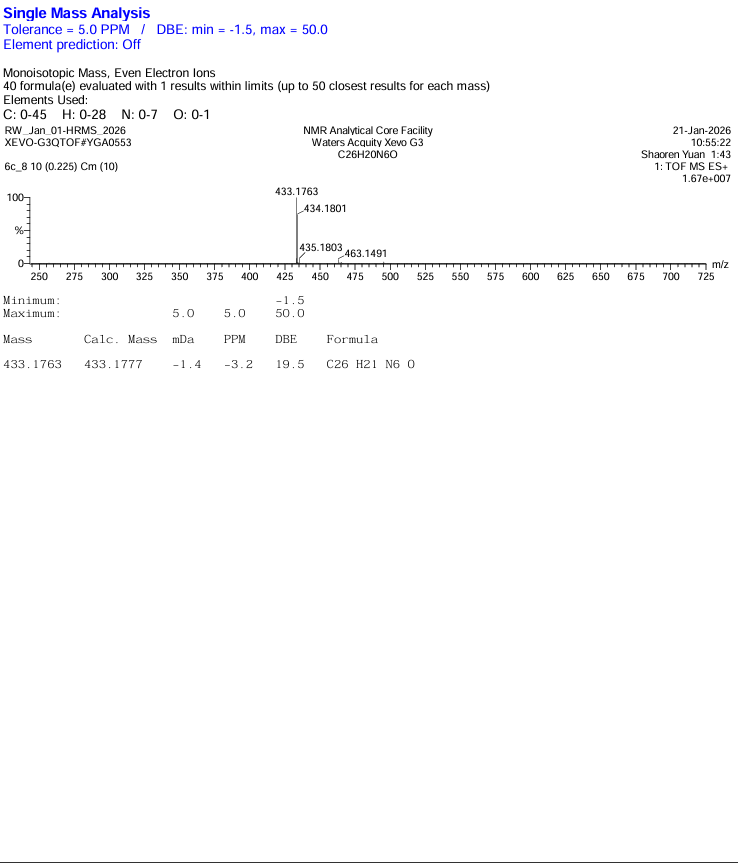

LC reports of compound **5** and **6**

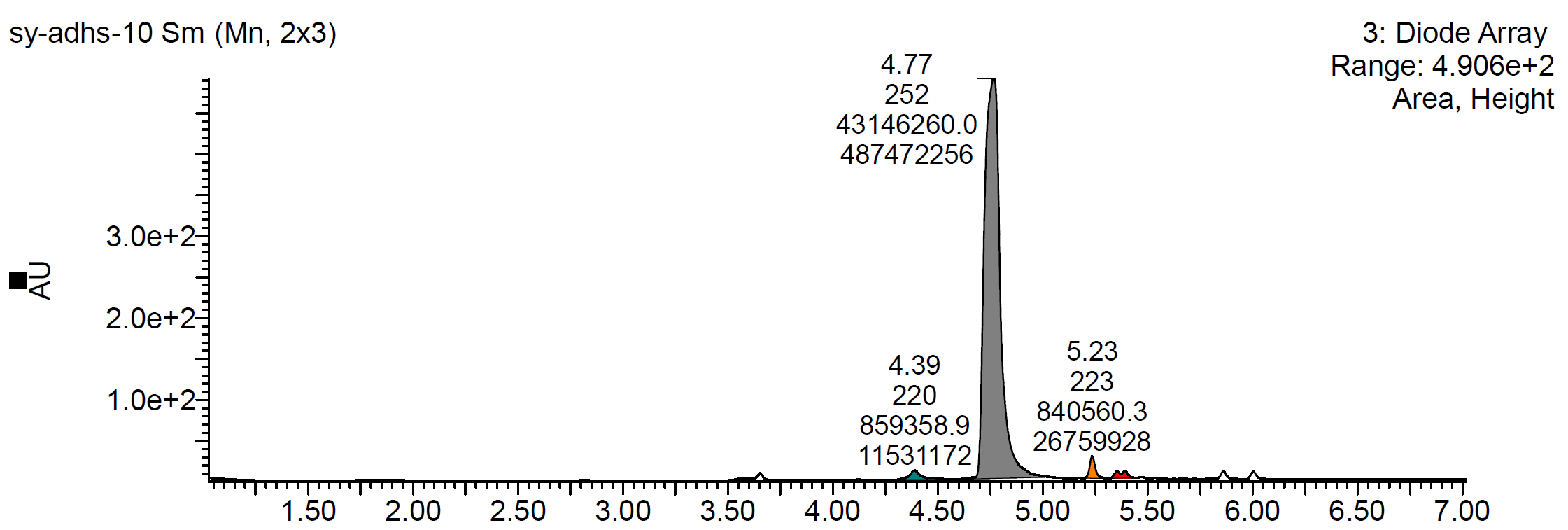
**5a:**

| Retention Time (min) | Area | Area% |
| --- | --- | --- |
| 4.39 | 859359 | 1.92% |
| 4.77 | 43146260 | 96.21% |
| 5.23 | 840560 | 1.87% |

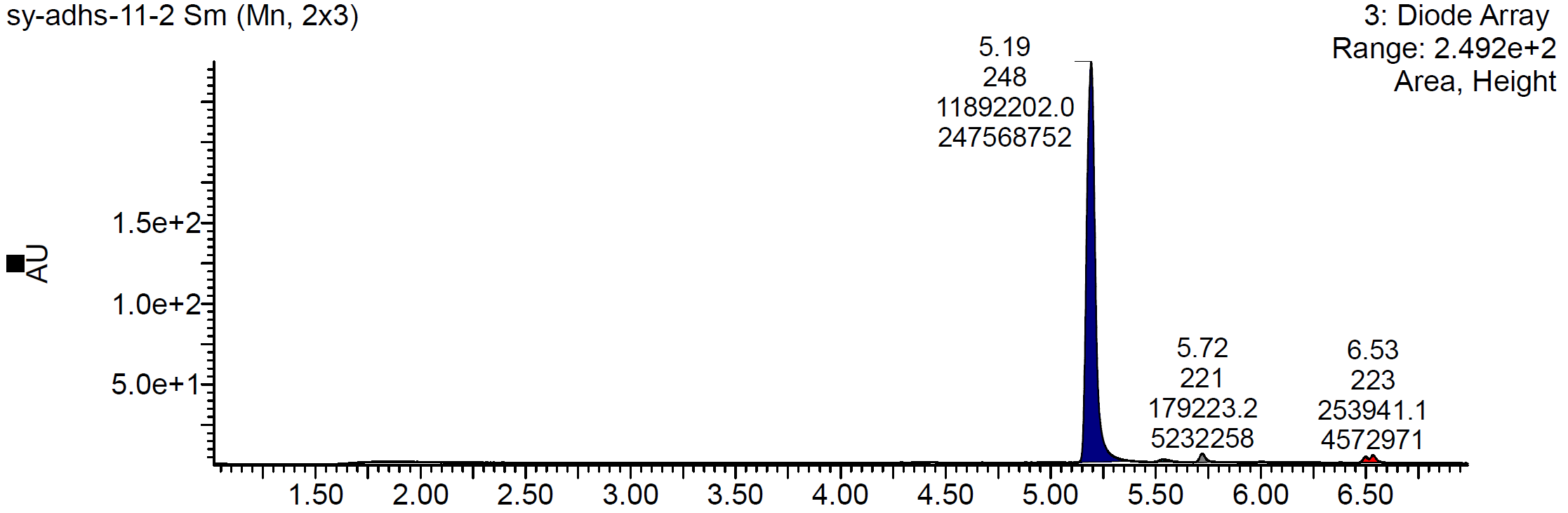

**5b**:

| Retention Time (min) | Area | Area% |
| --- | --- | --- |
| 5.19 | 11892202 | 96.49% |
| 5.72 | 179223 | 1.45% |
| 6.53 | 253941 | 2.06% |

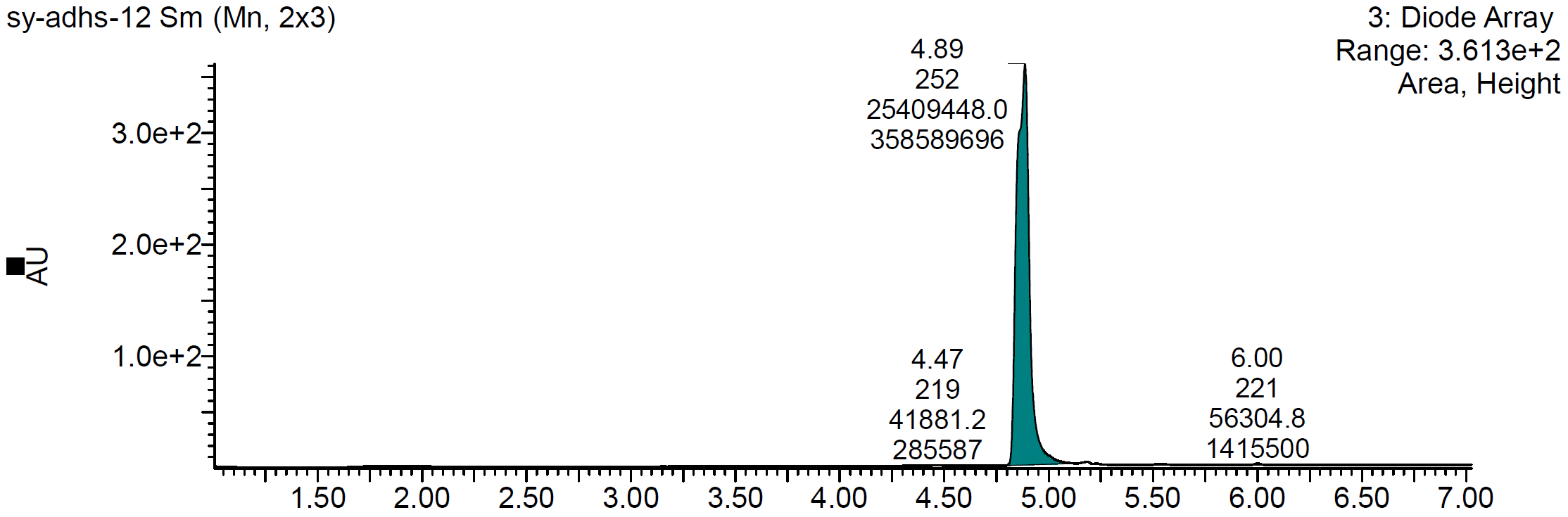
**6b:**

| Retention Time (min) | Area | Area% |
| --- | --- | --- |
| 4.47 | 41881 | 0.16 |
| 4.89 | 25409448 | 99.62 |
| 6.00 | 56305 | 0.22 |

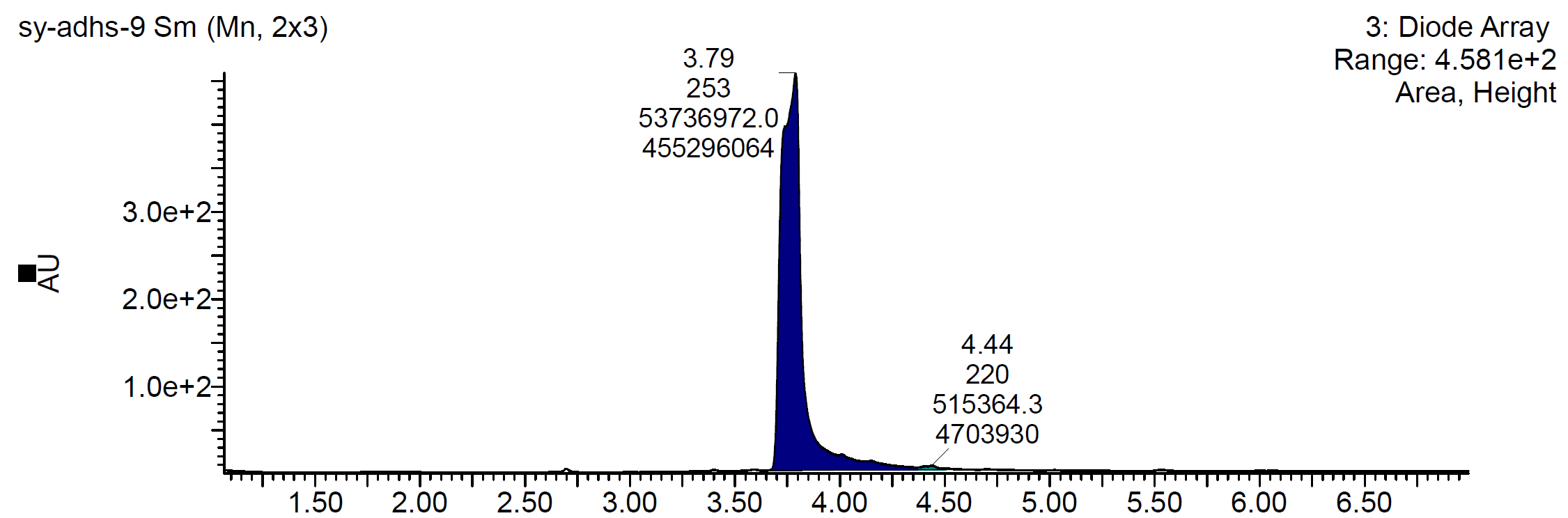

**5c:**

| Retention Time (min) | Area | Area% |
| --- | --- | --- |
| 3.79 | 53736972 | 99.05 |
| 4.44 | 515364 | 0.05 |

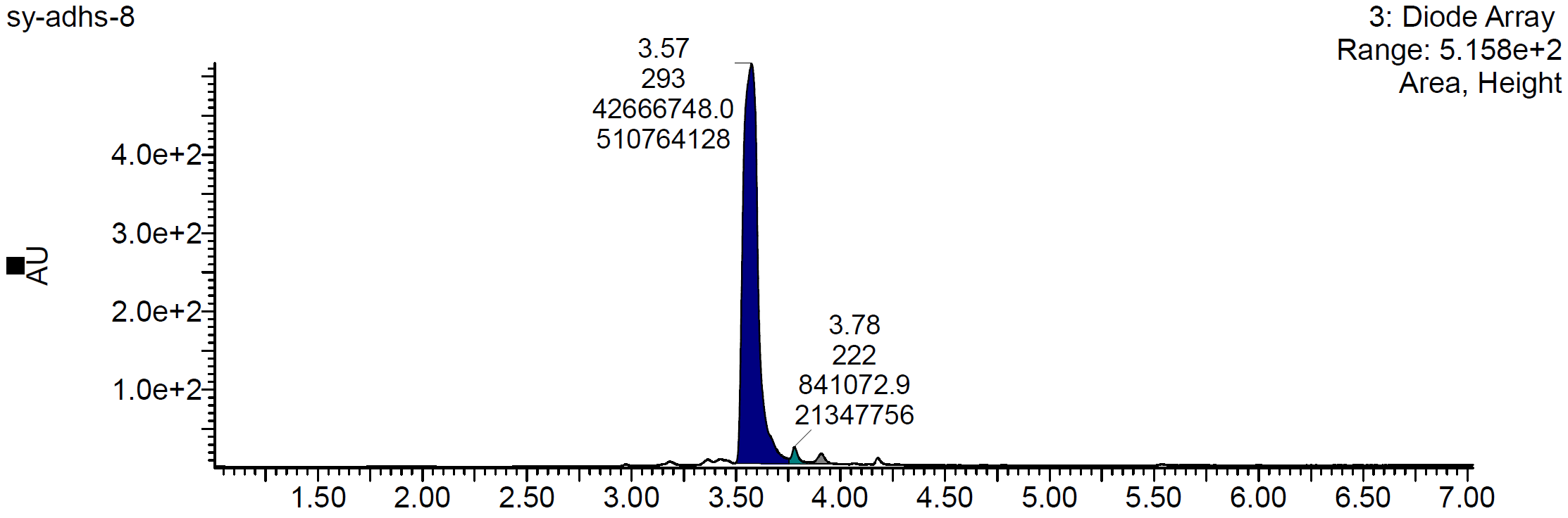
**6c:**

| Retention Time (min) | Area | Area% |
| --- | --- | --- |
| 3.57 | 42666748 | 96.74 |
| 3.78 | 841073 | 1.91 |
| 3.91 | 598282 | 1.35 |
